# Structural Basis of Membrane Potential Coupled Vectorial CO_2_ Hydration by the DAB2 Complex in Chemolithoautotrophs

**DOI:** 10.64898/2026.03.13.711513

**Authors:** Yat Kei Lo, Michael Seletskiy, Stefan Bohn, Darja Deobald, Timo Glatter, Sven T. Stripp, Jan M. Schuller

**Affiliations:** CryoEM of Molecular Machines, SYNMIKRO Research Center and Department of Chemistry, Philipps University of Marburg, Marburg, Germany; Institute of Structural Biology, Helmholtz Munich, Ingolstädter Landstraße, 85764 Neuherberg, Germany; Molecular Environmental Biotechnology, Helmholtz Centre for Environmental Research (UFZ), Permoserstraße 15, 04318 Leipzig, Germany; Facility for Mass Spectrometry and Proteomics, Max Planck Institute for Terrestrial Microbiology, Karl-von-Frisch-Str. 10, 35043 Marburg, Germany; Spectroscopy & Biocatalysis, Insitute of Chemistry, University of Potsdam, Karl-Liebknechts-Strasse 24/25, 14476 Potsdam, Germany

## Abstract

The fixation of dissolved inorganic carbon (DIC) such as CO_2_ and bicarbonate by autotrophic microorganisms is fundamental to the global primary production. Many autotrophs depend on a diversity of CO_2_-concentrating mechanisms (CCMs) to overcome the inefficiency of ribulose-1,5-bisphosphate carboxylase/oxygenase (RuBisCO) and the limited supply of DIC. While cyanobacterial CCMs are well characterized, analogous systems in chemolithoautotrophs, specifically active DIC uptake systems have long been overlooked. Here, we present the first cryo-EM structural analysis of DAB2, an essential membrane-associated protein complex for CO_2_ uptake in *Halothiobacillus neapolitanus*. The cytoplasmic subunit DabA2 displays a β-carbonic anhydrase-like fold including a zinc ion, while the transmembrane subunit DabB2 resembles the proton-conducting subunits of respiratory Complex I. Purified DAB2 binds CO_2_ independent of proton motive force (PMF) however, did not spontaneously hydrate CO_2_. This suggests that CO_2_ hydration is PMF-dependent and may involve a gating mechanism. Structural analysis reveals an unconventional deeply buried active site only accessible via gated substrate tunnels, implying that substrate access, product release, and catalytic activation are tightly regulated. A unique transmembrane helix of DabA2 constitutes part of the proton conduction pathway and potentially couples proton translocation to enzymatic turnover. These features define a vectorial CO_2_ hydration mechanism that prohibits reverse bicarbonate dehydration and requires a proton gradient to initiate catalytic turnover. Our findings establish DAB2 as a prototype of a previously unrecognized family of PMF-driven carbonic anhydrases, elucidating a novel strategy for CO_2_ capture in non-photosynthetic autotrophs and expanding the mechanistic landscape of bacterial CCMs.

## Introduction

The autotrophic carbon fixation by microorganisms is essential for global primary production and forms the basis of food chains in a variety of ecosystems, ranging from oceanic photic zones to hydrothermal vents. A key part of this process is the Calvin–Benson–Bassham (CBB) cycle, in which ribulose-1,5-bisphosphate carboxylase/oxygenase (RuBisCO) catalyzes the incorporation of CO_2_ into organic molecules. Despite of its importance, RuBisCO has a low turnover rate and is susceptible to competitive inhibition by O_2_, which results in a loss of fixed carbon and ATP through photorespiration^1^. Furthermore, in many natural environments, the availability of dissolved inorganic carbon (DIC), the collective pool of aqueous CO_2_, bicarbonate (HCO₃⁻) and carbonate (CO₃²⁻), is often restricted due to slow gas exchange, sensitivity to environmental pH and temperature^2,3^. These challenges have driven the widespread evolution of CO_2_-concentrating mechanisms (CCMs) – protein complexes and microcompartments that accumulate cytoplasmic HCO₃⁻and elevate the local CO_2_ concentration around RuBisCO to enhance carbon fixation^4^.

Cyanobacteria are the dominant oxygenic phototrophs in many aquatic ecosystems and have developed highly effective CCMs that serve as a benchmark for understanding microbial carbon acquisition^4–6^. These systems rely on two principal components: (I) energy-coupled inorganic carbon uptake systems that directly import HCO₃⁻ or indirectly via hydrating cytoplasmic CO_2_ to HCO₃⁻ against the concentration gradient, and (II) proteinaceous microcompartments, known as carboxysomes which encapsulate RuBisCO alongside carbonic anhydrase (CA), the later providing the substrate for the former based on the DIC pool^7^. A variety of cyanobacterial transporters with different affinities and capacities has been investigated, including the ATP-binding cassette (ABC) type HCO₃⁻ transporter BCT1^8^, the Na⁺-dependent, high-affinity symporter SbtA^9^, and the low-affinity, high-flux BicA transporter^10^. Additionally, some cyanobacteria express vectorial carbonic anhydrases (vCAs) such as the NDH-1MS and NDH-1MS’ complexes that function by coupling ferredoxin oxidation to CO_2_ hydration at the thylakoid membrane^11,12^. Together, these systems enable cyanobacteria to establish a high cytoplasmic bicarbonate concentrations and achieve carboxysomal CO_2_ levels that are several orders of magnitude higher than the extracellular levels, thereby saturating RuBisCO and minimizing photorespiration^4^.

By contrast, active DIC uptake systems of chemolithoautotrophs, organisms that fix CO_2_ using energy derived from inorganic redox reactions, remain poorly understood. Many such species inhabit environments with low and fluctuating DIC concentrations, including hydrothermal vents and sulfide-rich sediments^2^, and are thus expected to have evolved specialized carbon uptake systems. A notable example is the DIC-accumulating complex (DAC) originally identified from deep-sea hydrothermal vent γ-proteobacterium *Thiomicrospira crunogena*^13^. The complex was co-transcribed by two genes (*Tcr_0853* and *Tcr_0854)* which were upregulated in response to DIC scarcity and demonstrated to be essential for growth under DIC limited condition^14^. Recently, a genome-wide barcoded transposon mutagenesis screen revealed novel variants of DAC in *Halothiobacillus neapolitanus*^15^. This species expressed a two-subunits DAC (DAB2) encoded by *hneap_0211* (DabA2) *and hneap_0212* (DabB2) similar to the Tcr_0853/0854 complex, as well as a three-subunits DAC (DAB1) encoded by *hneap_0907* (DabA1), *hneap_0909* (DabB1), and an additional small hypothetical transmembrane protein (*hneap_0908)*. While the cytoplasmic subunits DabA1 and DabA2 were classified as *probable inorganic carbon transporter subunits* (Pfam PF10070; formerly as domain of unknown function DUF2309), DabA2 was predicted to harbor a zinc-dependent active site similar to that found in β-carbonic anhydrase (β-CA) or ζ-carbonic anhydrase (ζ-CA). The transmembrane subunit DabB1 and DabB2 were homologues of NADH:quinone oxidoreductases and Mrp antiporters (Pfam PF00361), suggesting a role in proton or sodium translocation. DabA2 and DabB2 formed a heterodimeric complex that was shown to accumulate intracellular DIC and rescue CA-deficient *E. coli* in a pH-independent manner^15^. Furthermore, i*n vivo* DIC-uptake experiments demonstrated that the complex utilized CO_2_ rather than bicarbonate as the main substrate, in which this activity was susceptible to CCCP uncoupling^16^. These observations hinted on the putative role of DAB2 as a proton or sodium motive force coupled vCA.

DACs were phylogenetically widespread and presented in at least 14 bacterial phyla, as well as in the Euryarchaeota^15^. Remarkably, homologues were not restricted to autotrophs but could also be found in heterotrophs and pathogenic species, including *Staphylococcus aureus*, *Vibrio cholerae*, and *Bacillus anthracis*. In contrast to the DAB system, the MpsAB complex, a DAC homologue expressed by *S. aureus*, was proposed to function as a Na⁺/HCO₃⁻ symporter and demonstrated to confer bacterial virulence, in addition to DIC accumulation, possibly in the form of HCO₃⁻ (ref.^17,18^). This illustrated the potential mechanistic diversity and physiological repurposing of this protein family.

Despite these functional insights, the molecular basis of DAC activity is still unclear. Key questions include how DACs are structurally organized, how CO_2_ is accumulated and converted, the source of energy and how it coupled to catalysis, and how the enzyme is regulated to prevent futile cycling. Notably, DAC differ fundamentally from canonical CAs in that their activity is not freely reversible and appears to depend strictly on membrane integrity and electrochemical gradients^15,16^.

Here, we present the first structural and mechanistic study of the DAB2 complex, a representative of the DAC transporter family underlying chemoautotrophic CO_2_-concentrating mechanisms. Our cryo-EM structures revealed a distinctive architecture possibly coupling proton transfer mediated by DabB2 to CO_2_ hydration catalyzed by the non-canonical vCA DabA2. Structural comparison with other CAs identified an unconventional active site in DabA2, connected by gated substrate tunnels. Additionally, DabA2 possesses a distinctive transmembrane helical extension that formed the putative proton conduit with DabB2. Infrared spectroscopy showed that the complex lacks catalytic activity in the absence of a membrane potential but exhibites strong CO_2_ binding. Our findings defined the DAB2 complex as a previously unrecognized class of proton motive force-driven carbonic anhydrases, thereby broadening the mechanistic diversity of bacterial CCMs.

## Results

### Cryo-EM analyses reveal DAB2 in multiple ligand-bound states

To unravel the molecular principle of DAC, we reconstituted the DAB2 complex from *H. neapolitanus* in lipid nanodiscs and determined its structure using cryo-EM single-particle analysis. Our initial attempts to prepare cryo-EM grids with the wild-type DAB2 complex were unsuccessful. Despite using mild detergents and synthetic polymers, the complex was highly unstable and prone to dissociation during sample preparation. To overcome this, we engineered a stabilized single-subunit variant by directly fusing the C-terminus of DabB2 to the N-terminus of DabA2, a construct we termed *Dab2*. This design mirrored the naturally occurring single-subunit DAC system found in *Acidimicrobium ferrooxidans*^16^ and had no significant impact on the activity of the complex (Supplementary Fig. 1a).

We captured *Dab2* in multiple functional states, providing unprecedented structural insights. By treating the protein with either ∼17 mM CO_2_ or 0.1 M bicarbonate (see Methods), we resolved its CO_2_-bound (*Dab2-CO_2_*, 2.8 Å) and bicarbonate-bound (*Dab2-HCO₃⁻*, 3.2 Å) conformations (Supplementary Fig. 2, 3). Unexpectedly, we also obtained a high-resolution structure of *Dab2* bound to CO_2_ (*Dab2-ambient*, 2.6 Å) from a separate sample prepared under atmospheric air (<430 ppm CO_2_, Supplementary Fig. 4). The exceptional quality of this density map, with a local resolution of 2.3 to 3.0 Å, allowed us to confidently model almost the entire protein, assign non-protein ligands with high accuracy, and identify multiple phospholipids associated with the transmembrane domain. Given the overall best resolution, *Dab2-ambient* served as the reference structure for subsequent analyses unless otherwise specified.

Consistent with previous observations^15^, our cryo-EM density maps revealed that *Dab2* assembled as a heterodimer, comprising the cytoplasmic subunit DabA2 positioned directly above the transmembrane subunit DabB2 (Fig. 1a). This structural arrangement was in excellent agreement with the molecular weight of approximately 200 kDa estimated by size-exclusion chromatography (Supplementary Fig. 1d). The overall structures were highly similar among samples prepared under different conditions with a root mean square deviation (RMSD) of 0.27 Å between *Dab2-ambient* and *Dab2-CO_2_*, as well as 0.47 between *Dab2-ambient* and *Dab2-HCO₃⁻*(Supplementary Fig. 6a).

**Figure 1.**
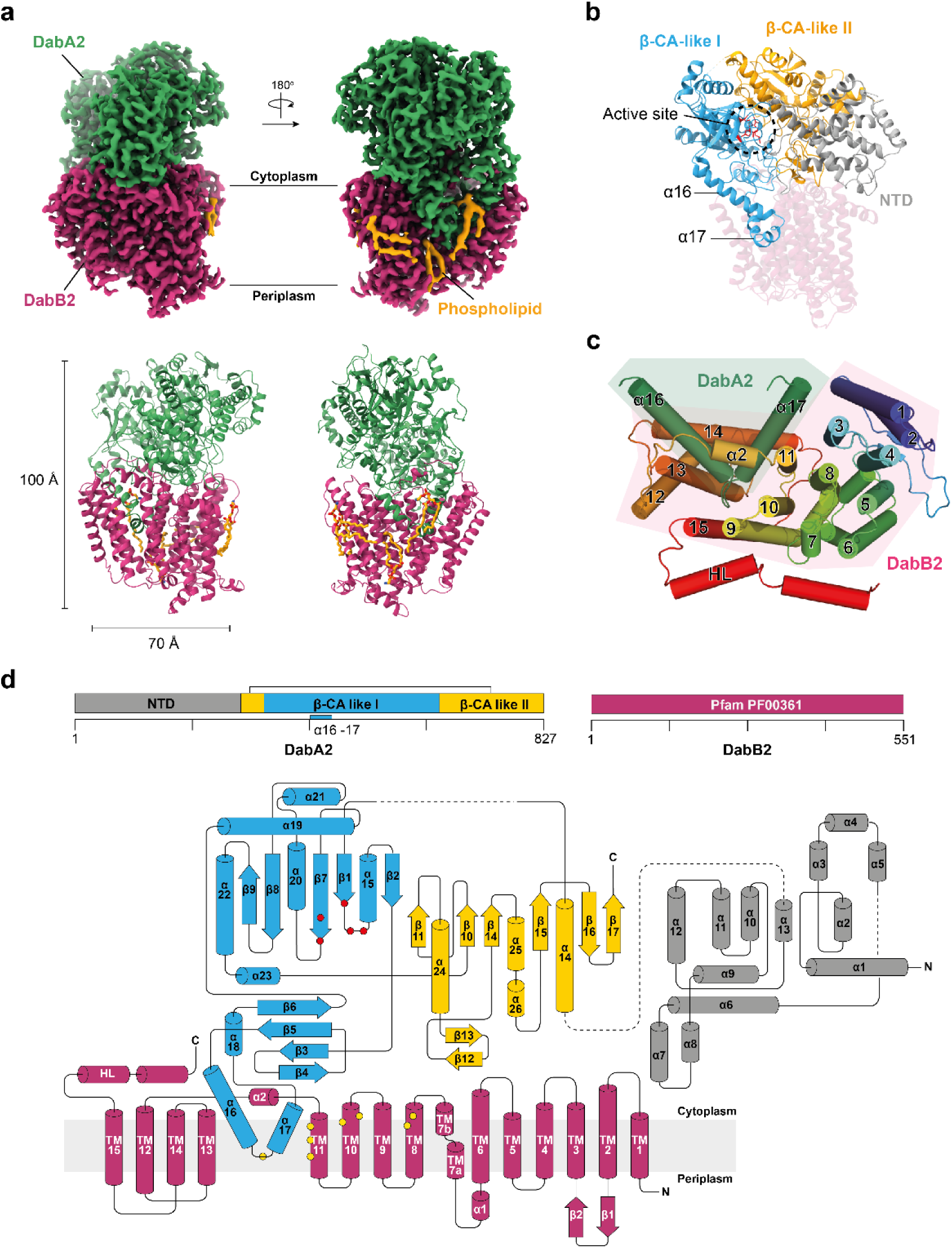
Overall structure of Dab2. **a** Cryo-EM density and structural model of the fusion protein complex *Dab2-Ambient* presented in front and back views. DabA2 and DabB2 are colored in green and magenta respectively. The density map revealed several phospholipids (in yellow) interacting with the membrane subunit, DabB2 and DabA2 transmembrane helices. **b** Domain architecture of DabA2. The catalytic core comprised of two β-CA-like domains (in cyan and yellow) and a N-terminal domain (NTD; in gray). Active site residues are highlighted in red. **c** Helices arrangement of DabB2 and DabA2 transmembrane “finger-like” motif (α16, α17) viewed from the cytoplasmic side. HL: lateral helix. **d** Topology of DAB2. Protein domains are colored according to **b**. Zinc-coordinating residues and putative proton transfer residues are depicted in red and yellow dots respectively. Dash lines indicate unresolved region, likely due to protein flexibility.

### DabA2 structurally mimics β-carbonic anhydrases

DabA2 was organized into three distinct domains: a helical N-terminal domain of 280 residues and two β-CA-like domains that form the catalytic core (Fig. 1b, d). Despite the structural resemblance, only five amino acids were strictly conserved between DabA2 and canonical β-CAs (Supplementary Fig. 7). The N-terminal domain comprised twelve intertwined α-helices and lacked any resemblance to other proteins, as confirmed by Foldseek^19^ and DALI^20^. Notably, this region was less conserved than the β-CA-like domains among DabA2 homologs (Supplementary Fig. 8), suggesting that its primary role is to maintain DabA2 proper folding rather than to participate directly in catalysis.

β-CAs are typically formed by two symmetrical Rossmann folds and function as a homo-dimer or homo-tetramer^21^. In contrast, the two β-CA-like domains in DabA2 displayed structural adaptations that broke this symmetry. The β-CA-like domain I (residues 341–655) featured an extended “finger-like” amphipathic helix–loop–helix motif (α16 and α17) which inserted deeply into the membrane bilayer, establishing interactions with DabB2 (Fig. 1b-d; Supplementary Fig. 9). Conversely, the β-CA-like domain II formed a discontinuous domain, comprising residues 281–336 and 656–827. It possessed an extended antiparallel β-sheet (β12 and β13), connecting the second and third β-sheets and form the cytoplasmic interaction interface (Fig. 1d). Collectively, this interface, together with the “finger-like” motif, accounts for a buried surface area of 4,700 Å² as calculated by the PISA server^22^. A dense network of hydrogen bonds and salt bridges further stabilized the interaction between DabA2 and DabB2, with hydrophobic contacts predominating between DabA2 helices α16/α17 and DabB2 transmembrane helices 11–14 (Supplementary Table 2, Supplementary Fig. 9). These intricate interfaces may contribute to the protein complex assembly and scaffold key residues for the catalytic function of the DAC transporter system (see below).

### DabA2 binds to CO_2_ and bicarbonate in a unique active site

DabA2 possessed a putative active site with a region of high electron density located within β-CA-like domain I, positioned near the interface of the two β-CA-like domains (Fig. 1b, 2a, Supplementary Fig. 6b-d). This correlated to the presence of one zinc ion per protein molecule, revealed using inductively coupled plasma mass spectrometry (ICP-MS; Supplementary Fig. 10). The catalytic zinc ion was coordinated by a Cys_2_His(H_2_O) motif comprised of Cys351, Cys539, and His524 (Fig. 2b), bearing resemblance to that observed in type I β-CAs or the “R-state” of type II β-CAs^21,23^. A zinc-bound water molecule or hydroxide ion was stabilized by the Asp353-Arg355 dyad. In β-CAs, the conserved Asp was proposed to mediate deprotonation of the zinc-bound water, a prerequisite for bicarbonate formation^24^. When we mutated the zinc-coordinating residues and the Asp-Arg dyad to alanine by site-directed mutagenesis, significantly impaired bacterial growth under CO_2_-limiting conditions was observed, underscoring their critical functional roles (Fig. 2e).The Ala substitutions had limited effect on the protein expression levels (Supplementary Fig. 11). Together with the structurally resolved water molecules hydrogen-bonded to conserved residues Ser356, His 585 and Asp590, Asp353 might serve a similar function in relaying the proton to the bulk solvent in DabA2 (Supplementary Fig. 12).

**Figure 2.**
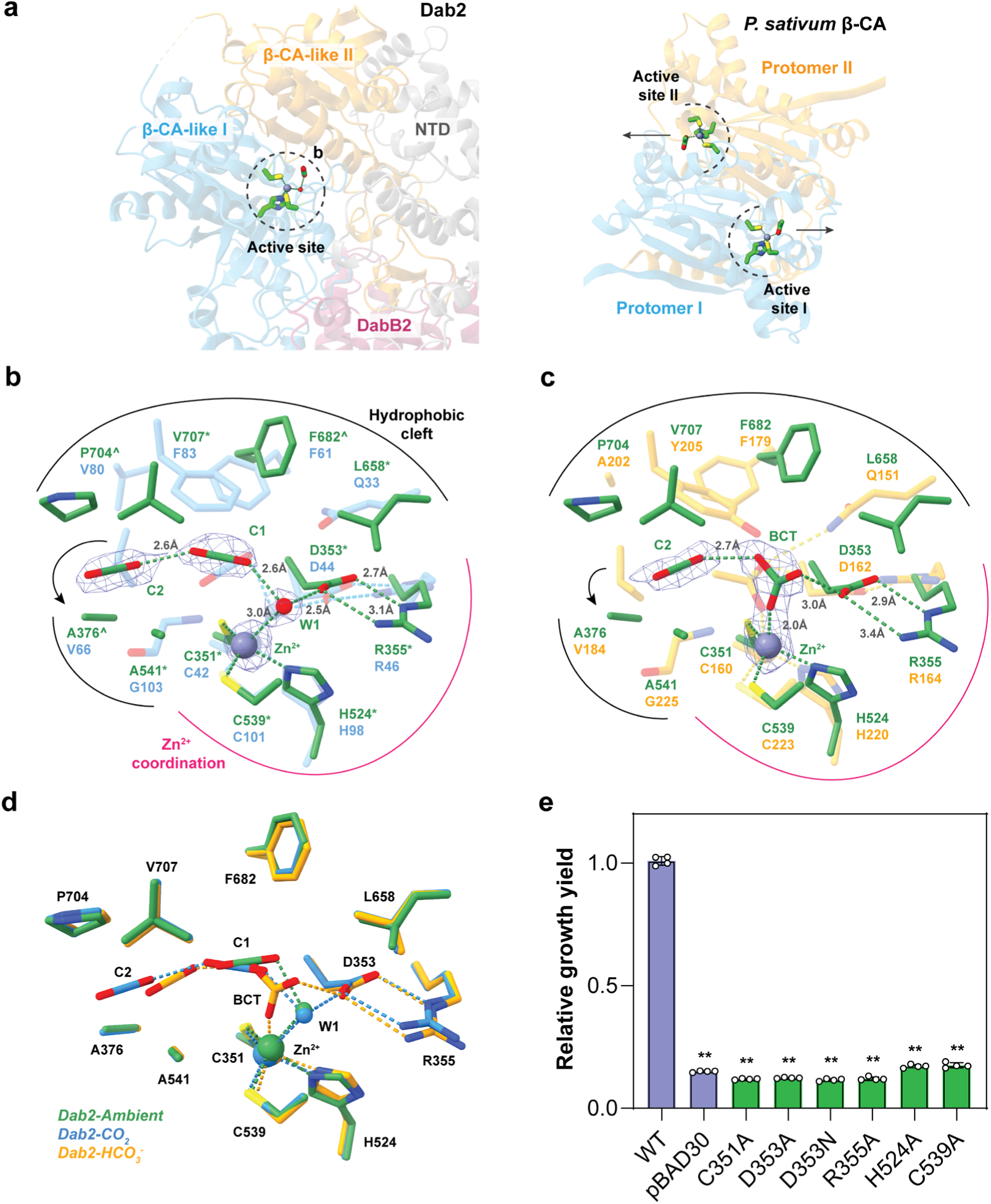
Active site architecture of DabA2 in CO_2_ and bicarbonate bound states. **a** Comparison of active site location in *Dab2* and *Pisum sativum* β-CA (PSCA; PDB 1EKJ). Arrows indicate immediate connection to the bulk solvent. **b** Superposition of *Pseudomonas aeruginosa* β-CA (PACA) in CO_2_-bound state (PDB 5BQ1; in blue) on *Dab2-CO_2_*, and **c** *PSCA* in acetate-bound state (in orange) on *Dab2-HCO_3_^-^* (right). Key molecular interactions are depicted by dashes. Residues strictly conserved or conserved at ≥ 90% sequence identity among DabA2 homologues are marked by asterisk (*) and circumflex (^) respectively. Arrows indicate reposition and substitution of β-CA conserved Val residue (Val66^PACA^ and Val184^PSCA^). Notably, the transition state stabilizing residue (Gln33^PACA^, Gln151^PSCA^) was substituted by Leu658 in DabA2. Densities of ligands are shown at 9.2 σ (*Dab2-CO_2_*) and 6.5 σ (*Dab2-HCO_3_^-^*). **d** Superposition of *Dab2* active site from *Dab2-ambient* (green), *Dab2-HCO_3_^-^*(blue), and *Dab2-CO_2_* (yellow) **e** Site-directed mutagenesis of Zn^2+^-coordinating residues. Bar heights and error bars represent means and standard deviations respectively (n = 4 biological replicates). “**” Indicates statistically significant difference compared to WT (P < 0.05) according to Holm-Bonferroni corrected two-tailed t-test. Bacterial growth yield is presented relative to that of wild-type (WT = 1.0).

Our structural analysis further revealed two elongated electron densities near the zinc ion, surrounded by hydrophobic residues in both *Dab2-CO_2_* and *Dab2-ambient*, which likely correspond to protein-bound CO_2_ molecules (Fig. 2b, Supplementary Fig. 6b-c). Their consistent appearance across independently prepared samples supported the assignment of these densities as specifically bound CO_2_. Conventional β-CAs typically accommodate only a single CO_2_ or bicarbonate molecule; however, the DabA2 active site appeared to bind two. The first CO_2_ molecule (C1) possibly formed a hydrogen bond with the zinc-bound water, suggesting the capture of a substrate-bound state. In β-CAs, the active site is enclosed by a loop and a conserved valine residue, precluding the binding of an additional CO_2_ molecule. Yet, in DabA2 the loop between β2 and β3 was displaced away together with the conserved valine replaced by Ala376, thereby expanding the active site cavity (Fig. 2b). This structural rearrangement permitted the accommodation of a second CO_2_ molecule (C2), stabilized by hydrophobic interactions within the active site cleft.

In *Dab2-HCO₃*, we identified a prominent triangular electron density adjacent to the zinc ion, likely representing a HCO₃⁻ bound at the active site (Fig. 2c). The structural similarity between *Dab2-HCO₃* and *Dab2-ambient* (RMSD = 0.47 Å), suggests substrates binding had minimal impact on the protein global conformation, as well as the active site architecture (Fig. 2d). While the HCO_3_ ^-^ bound state might be captured during catalytic turnover or due to binding of added ion, we cannot exclude the possibility that this density arises from the non-enzymatic CO_2_ hydration due to an increased pH from 7.5 to 7.8 upon addition of NaHCO₃. Even if this was the case, the observed bicarbonate still occupied the active site in a manner comparable to catalytically active β-CAs^25^, suggesting a conserved mode of substrate coordination.

To validate the binding of supernumerary CO_2_ within the *Dab2* protein complex independent of structural analysis, we employed attenuated total reflectance (ATR) FTIR spectroscopy^26^. This is a robust method for detecting CO_2_/HCO₃⁻ conversion and CO_2_ protein binding^27–29^. A Dab2 protein film was formed under inert carrier gas (N_2_) before the reaction was started by changing the atmosphere *in situ* to 10% CO_2_, and time-resolved “CO_2_-minus-N_2_” difference spectra were calculated by subtracting the N_2_ background spectrum from all spectra recorded under 10% CO_2_ (Fig. 3a, b; see Supplementary Fig. 13a for the complete datasets). Bands at 1614 cm⁻¹, 1360 cm⁻¹, and 1302 cm⁻¹ were assigned to the stretching (ν_2_, ν₃) and bending (ν₄) modes of HCO₃⁻ (ref.^29^). These bands appeared positive in the spectra of *E. coli* β-CA (ECCA), equivalent to a fast increase of HCO₃⁻ upon enzymatic hydration of CO_2_. A feature at higher frequencies was fitted with bands at 2341 and 2337 cm^-1^, indicative of CO_2_ in aqueous solution and CO_2_ bound to protein, respectively^27^. This shift relates to differences in proticity and can be mimicked by comparing the IR spectrum of CO_2_ in H_2_O or DMSO (Supplementary Fig. 14). Overall similar results were observed for BSA, which were utilized as a protein standard to probe “non-catalytic” CO_2_ hydration. The HCO₃⁻ formation in BSA was significantly slower than ECCA (Fig. 3b, c), and a similar response was observed for *Dab2*, hinting at non-catalytic CO_2_ hydration with both BSA and Dab2. In contrast, the CO_2_ feature of *Dab2* was dominated by the 2337 cm⁻¹ band, approximately ten times stronger than the aqueous CO_2_ signature observed for BSA and ECCA (Fig. 3a). This indicated that membrane potential is not essential for CO_2_-binding, even though the protein complex itself did not spontaneously catalyze CO_2_ hydration under these conditions.

**Figure 3.**
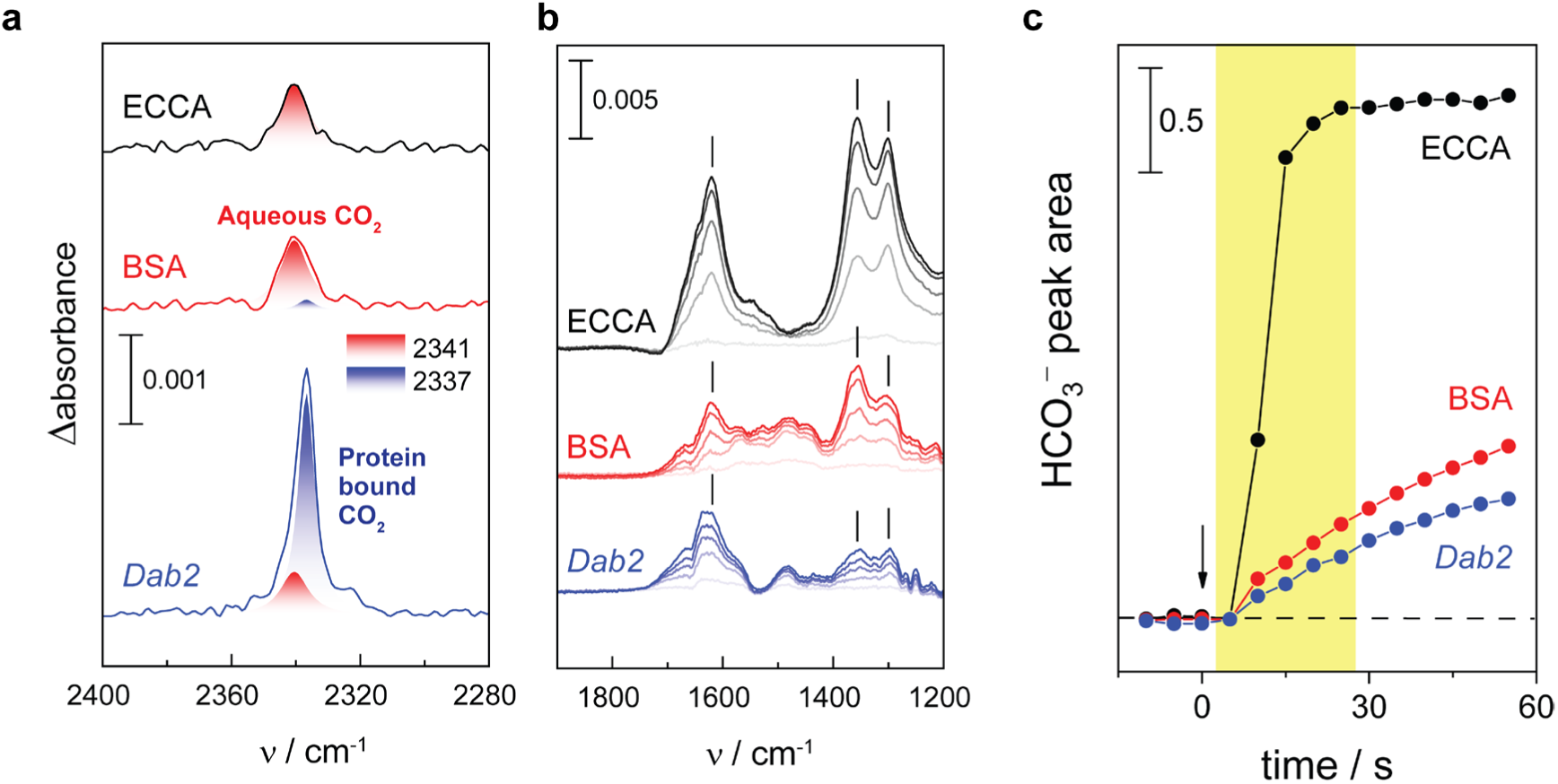
Comparison of CO_2_ binding and bicarbonate formation. Time-resolved “CO_2_-minus-N_2_” ATR FTIR difference spectra for *E. coli* β-CA (ECCA, black), BSA (red), and *Dab2* (blue). **a** After 25 s in the presence of 10% gaseous CO_2_, bands at 2341 and 2337 cm^-1^ indicate the presence of dissolved CO_2_ or protein-bound CO_2_, respectively. **b** At lower frequencies, the increasingly more positive bands at 1614, 1360, and 1302 cm⁻¹ revealed HCO_3_^−^ formation. All spectra run from light color (t = 5 s) to full color (t = 25 s). **c** Kinetics of CO_2_ hydration with ECCA, BSA, and *Dab2*. The arrow marks the switch from 100% N_2_ to 90% N_2_ and 10% CO_2_(t_0_). Spectra in panel **b** relate to the time frame between 5–25 s highlighted in panel **c**.

### DabA2 lacks the canonical residue for stabilizing the transition state

To understand why DabA2 did not readily catalyze CO_2_ hydration, we performed a detailed inspection of its active site and compared it with canonical β-CAs. In β-CAs, a conserved glutamine or histidine residue was suggested to stabilize the transition state during catalysis by hydrogen-bonding to the zinc-bound intermediate^25,30^, in which substitution of the conserved Gln/His impaired CA activity^31,32^. An exception is observed in *Mycobacterium tuberculosis* β-CA Rv1284, which, despite lacking any equivalent charged or polar residues, retains CA activity. Its high-resolution structure revealed a water molecule occupying the position of the canonical Gln/His sidechain that may serve as an alternative hydrogen bond donor^33^. In stark contrast, DabA2 lacks any analogous residues, as the conserved Gln/His was replaced by a hydrophobic residue (Leu658), which is incapable of fulfilling this role and does not leave room for a water molecule (Fig. 2b). Notably, this substitution was strictly conserved among 150 DabA2 homologues. To assess whether the absence of a canonical hydrogen-bonding residue renders DAB2 dependent on membrane potential, we substituted Leu658 with glutamine to mimic a canonical β-CA. Although this variant supported bacterial growth under CO_2_-limiting conditions, it failed to catalyze CO_2_ hydration *in vitro* (Supplementary Fig. 15), implying that catalysis remains contingent upon a membrane potential. Unexpectedly, replacing Leu658 with glycine, alanine, or asparagine also did not impair DAB2 *in vivo* activity (Supplementary Fig. 15b). These observations demonstrated that Leu658 was not essential for the function despite being strictly conserved. DabA2 might employ a novel mechanism for CO_2_ hydration, independent of the canonical Gln or His residue. Overall, the lack of spontaneous CA activity likely arises from a more intricate structural configuration that extends beyond a mere single residue substitution at the active site.

### The deeply buried active site in DabA2 requires a CO_2_/HCO_3_^-^ tunnel

The active site of DabA2 was embedded within the protein core, in contrast to the more surface-exposed active sites of conventional β-CAs (Fig. 2a). This structural arrangement mandates a specialized adaptation—a tunnel system that facilitates efficient diffusion of CO_2_ and HCO₃⁻ to and from the active site^34,35^. Using CAVER^36^, we identified two putative tunnels (T1, T2) extending from the active site in opposite directions, leading to the bulk solvent (Fig. 4a). The first tunnel was lined with hydrophobic residues with a bottleneck radius of 1.3 Å and branched into two entrances (T1a, T1b). This bottleneck was defined by conserved residues Ala376, Phe378, Pro704, and Ala703 (Fig. 4b, c). In addition, tunnels T1b traversed a second bottleneck of similar dimensions, formed by Phe375, His684, Val696, and Ile700, located immediately downstream. Remarkably, density maps of both *Dab2-ambient* and *Dab2-CO_2_* revealed several probable CO_2_ molecules (designated C3 to C5) within and near the tunnel, stabilized by hydrophobic residues (Fig. 4a, Supplementary Fig. 16a, b). This observation was in excellent agreement with the IR signature of protein-bound CO_2_ (Fig. 3a). Although the predicted bottleneck is slightly narrower than the radius of CO_2_ molecule (∼1.6 Å), protein flexibility may permit CO_2_ diffusion without significant conformational changes.

**Figure 4.**
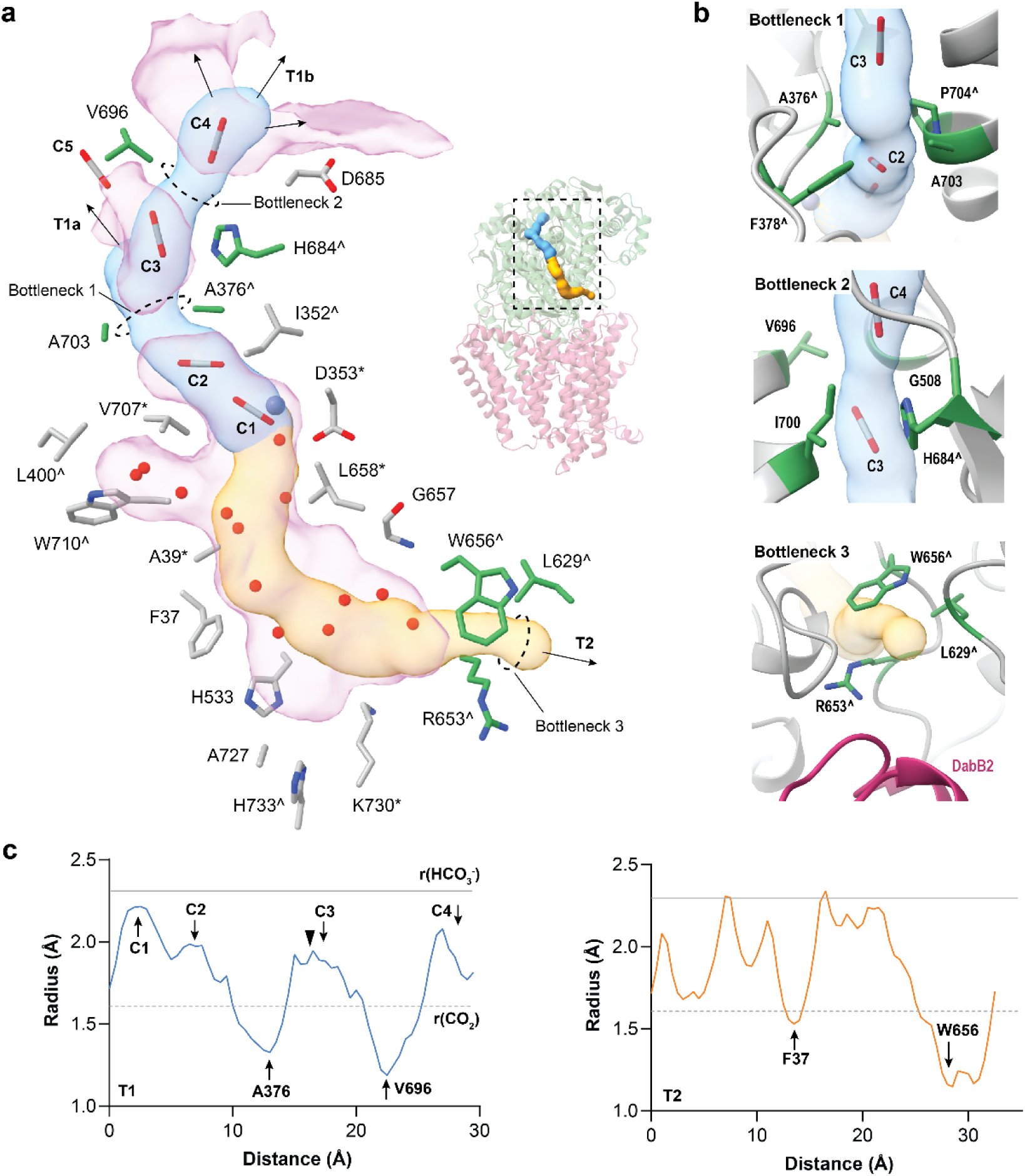
Prediction of tunnels connecting the active site. **a** CAVER 3 predicted 2 tunnels (T1, in blue; T2, in orange), connecting the active site to the bulk solvent. T1 was populated with four potential CO_2_ molecules (C1-C4), and branched into two openings (T1a, T1b). Another potential CO_2_ molecule (C5) resided near the entrance of T1a. T2 was hydrated by a number of structurally resolved water molecules. Residues lining the tunnel are shown in gray. Bottleneck residues are shown in green. Arrows indicate connection to the bulk solvent. Pink blobs represent cavity calculated by KVFinder^61^. Residues strictly conserved or conserved at ≥ 90% sequence identity among DabA2 homologues are marked by asterisk (*) and circumflex (^) respectively. See Supplementary Fig. 16 for the detailed ligand coordination. **b** Zoomed-in view of the tunnel bottlenecks. **c** Radius along the predicted tunnels, extending from the zinc ion (blue line: T1, orange line: T2). Gray dotted line and solid line indicate the radius of CO_2_ and HCO_3_^-^ respectively. Arrows mark position of bottlenecks and CO_2_ molecules. Black triangle indicates the T1a opening.

*Dab2* possesses a second putative tunnel (T2) that connects the active site to the bulk solvent near the DabA2–DabB2 interface, passing through a large cavity (Fig. 4a). Although primarily scaffolded by hydrophobic residues, several water molecules were found in T2 hydrogen-bonded to backbone amides, which could offer a more favorable, polar route for HCO₃⁻ egress. This tunnel featured a bottleneck radius of 1.1 Å, gated by conserved residues Arg653, Trp656, and Leu629 (Fig. 4b, c). Given both bottlenecks are significantly narrower than the radius of HCO₃⁻ (∼2.3 Å), it suggests that *Dab2* must undergo substantial conformational rearrangements to facilitate product release via either tunnel. Collectively, these observations highlighted the intricate structural adaptations that govern substrate access and product release in DabA2.

### DabB2 and DabA2 form a putative proton conduit

DabB2 was classified as a proton-conducting membrane transporter (Pfam family PF00361) based on sequence similarity^15^. In agreement with this classification, our structure revealed that DabB2 shared a partial structural homology with NuoL, the distal proton-pumping subunit of *E. coli* respiratory Complex I (Fig.5a, b, Supplementary Fig. 17). Both NuoL and DabB2 features 15 transmembrane helixes, in contrast to 14 in NuoM and NuoN. Moreover, DabB2 possessed a short axial helix (HL), partly resembled that of NuoL (residues 499-527; Fig. 1d, 5a) which is absence in NuoM/N. Notably, DabB2 transmembrane segments TM1–11 aligned closely with those of NuoL, yielding an RMSD as small as 1.05 Å over 235 Cα atoms (residues 82-210, 313-354, 221-238, 251-296). However, subtle differences in helix positioning, combined with the integration of DabA2 extended “finger-like” motif, yielded an overall architecture distinct from NuoL.

**Figure 5.**
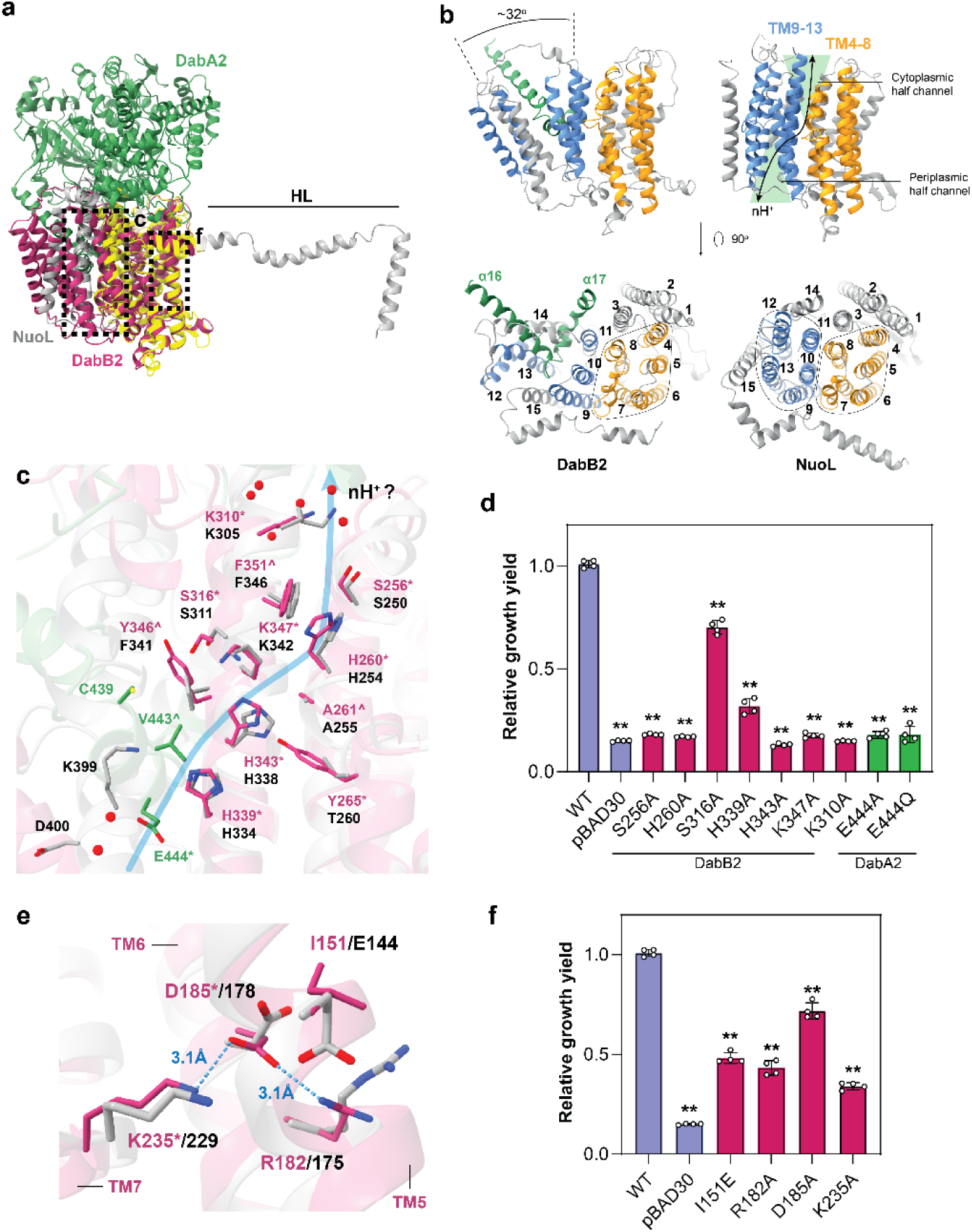
Similarities between DabB2 and NuoL revealed a putative proton pathway. **a** Superposition of NuoL (PDB 7P62; in gray) on DabB2. Region aligned to DabB2 is colored in yellow. Notice that DabB2 lacked the long axial helix (HL) found in Complex I proton-pumping subunits. **b** Topological comparison between DabB2 and NuoL. DabB2 antiparallel helix bundles (in blue and orange) were rearranged to accommodate DabA2 “finger-like” motif (α16, 17; in green). Arrow indicates the proposed NuoL proton transfer pathway. **c** Key residues along the proton pathway were conserved between DabB2 (in magenta) and NuoL (in gray), suggesting *Dab2* might conduct protons in a similar way (depicted by blue line). DabA2 is colored in green. The putative pathway opening was hydrated by several structurally resolved water (red spheres). Residues strictly conserved among DabA2 or DabB2 homologues are marked by asterisk (*). Residues conserved at ≥ 90% sequence identity among homologues are marked by circumflex (^). See Supplementary Fig. 19 for the detailed water coordination. **d** Substitutions of the polar and charged residues indicated their importance for DAB2 activity. **e** Comparison between DabB2 and NuoL terminal regulatory ion-pairs. Residues of ≥80 % identity between DabB2 homologues are marked by asterisk (*). Blue dashes depict interactions between DabB2 ion-pairs. **f** Interruptions of these ion-pairs reduced DAB2 activity. **d**, **f** Bar heights and error bars represent means and standard deviations, respectively (n = 4 biological replicates). “**” Indicates statistically significant difference compared to WT (P < 0.05) according to Holm-Bonferroni corrected two-tailed t-test.

NuoL has been proposed to transport protons via two antiparallel helix bundles (TM4–8 and TM9–13) that form an S-shaped, discontinuous water channel lined with polar or charged residues and interspersed by a hydrophobic barrier at the membrane’s midplane^37–39^. In DabB2, TM4–11, including the segmented helix TM7a and TM7b closely resembled NuoL’s cytoplasmic half-channel (Fig. 5b). Intriguingly, residues responsible for proton transfer in NuoL were also conserved in DabB2, in which alanine substitutions at these positions significantly impair DAB2 activity (Figs. 5c, d). These observations suggest that DabB2 TM4–11 likely function analogously to NuoL’s proton channel.

The major structural divergence was observed in TM12–14. In NuoL, TM12 and TM13 are tightly associated with TM9–11, whereas in DabB2 these helices were tilted outward, repositioning TM14, and TM15 (Fig. 5b). This arrangement exposed the hydrophobic core to the cytoplasm and facilitated interaction with DabA2’s “finger-like” motif (Fig. 1d, 5b). A conserved glutamate (Glu444) within this motif took the place of Lys399 and Asp400 in NuoL’s broken helix TM12b (Fig. 5c) that forms the periplasmic half-channel entrance^38,40^. Lys399 and Asp400 are involved in the oxidoreductase activity and proton pumping of complex I^41^, correspondingly, substituting Glu444 with alanine or glutamine abolished DAB2 activity *in vivo* (Fig. 5d). This indicates that Glu444 does not merely function as a water-bonding residues but it might directly take part in the proton transfer through protonation and deprotonation. While DabB2 lacked a helix bundle equivalent to TM9–13 in NuoL, the integration of DabA2’s “finger-like” motif with DabB2 TM12 and TM13 formed a probable periplasmic half-channel for the proton transfer. This distinctive configuration was highly conserved among DabB proteins and likely constitutes a critical coupling point for mediating DAB2’s proton-coupled vectorial carbonic anhydrase activity.

Interestingly, DabB2 possessed a structural adaptation that may enhance its proton transfer efficiency. In NuoL, two conserved ion pairs (Lys229/Asp178 and Arg175/Glu144) located near the transmembrane subunits interface have been proposed to alternate between an ‘open-state’ and ‘close-state’ conformation in response to electron transfer across the peripheral arm of Complex I, thereby modulate lateral proton transfer between the two half channels^42,43^. While the first ion pair was retained in DabB2 (Lys235/Asp185), the Glu residue necessary for the ‘close-state’ conformation was substituted by Ile151 (Fig. 5e). This adaptation dissociated the second ion pair and possibly stabilizes DabB2 in the ‘open-state’ conformation, reducing the energy barrier for proton transfer^43^. To probe the function of these ion pairs, we disrupted their interactions by individually replacing them with alanine by site-directed mutagenesis. All variants could complement CA-deficient *E. coli* strains; however, their growth yield was reduced by 30% to 60% (Fig. 5e). Furthermore, reintroducing a Glu residue in position 151 – which should enable alternation between both conformations, reduced the growth yield by half. These observations supported that the ion pairs modification is required for the optimal coupled catalysis.

While our structural analysis suggests that DAB2 maybe coupled to proton transfer, a homologue, the MpsAB complex has been argued to harness sodium gradient instead^17,44^. To clarify this ambiguity, we performed a comparable complementation assay as previously reported in sodium transporters deficient strain (Δ*nhaAB*)^44^, expressing DAB2. However, unlike MpsAB, DAB2 could not rescue the mutant strain under sodium stress (Supplementary Fig. 18a). Similarly, DAB2 retained full activity when cells were cultivated in the absence of sodium (Supplementary Fig. 18b). These observations imply sodium is probably not required for DAB2 and that the protein complex appears to transport protons exclusively.

## Discussion

In this study we present the first structural characterization of the DAB2 complex, a representative of membrane potential-dependent CO_2_ hydration systems found in chemolithoautotrophic bacteria. The structures revealed a unique architecture which differs mechanistically from the cyanobacterial CCMs that have been characterized to date. This has allowed us to define a previously unrecognized family of vectorial carbonic anhydrases (vCAs) which couple unidirectional CO_2_ hydration to transmembrane proton flux.

Our analysis pinpointed several distinctive features that intricately link carbonic anhydrase activity to proton transfer. We obtained structures of DAB2 in lipid nanodiscs in CO_2_ and bicarbonate-bound states. DabA2 adopts a protein fold homologous to dimeric β-class CA, but its active site architecture diverges substantially. Specifically, the residue that stabilizes the transition state in canonical carbonic anhydrases is replaced by Leu658 in DabA2, which disrupts critical hydrogen bonding and potentially contributes to the enzyme’s low basal activity in the absence of proton transfer. Additionally, the active site is buried with the protein core and only accessible via narrow, gated tunnels. This likely impose kinetic barriers to CO_2_ entry and bicarbonate release. These features suggest a latent catalytic core that is only activated upon proton-driven conformational rearrangement.

DabB2, a membrane-integral subunit, is central to this activation mechanism. It exhibits significant architectural similarity to the antiporter-like subunits of respiratory Complex I, particularly NuoL. Unlike canonical antiporters, however, DabB2 lacks the key ion-pair motifs required for regulating directional proton-pumping, and its structure is modified by the insertion of a long helical extension from DabA2. This ‘finger-like’ transmembrane helix appears to integrate into DabB2, likely contributing to the formation of a periplasmic half-channel for proton conduction. Its placement at the interface of the proton translocation machinery suggests that DabA2 has a dual role as both a structural modulator of proton flux and a sensor of local protonation dynamics. Mutational analysis of conserved residues along the putative proton conduction pathway supports a model in which proton-driven conformational changes initiate catalysis. Together with comparisons to the MpsAB complex, these findings argue against sodium-dependent coupling and instead support PMF-driven activation of DAB2.

Taken together, we hypothesize a PMF-driven vectorial CO_2_ hydration mechanism (Fig. 6). In the absence of a proton flux, CO_2_ and water can enter the active site via the substrate tunnel. However, HCO₃⁻ binding is prevented due to steric hindrance, thus hindering reverse dehydration. A zinc-bound water deprotonates to form a hydroxide ion, with the liberated proton potentially being conducted to the bulk solvent likely via the conserved residue Asp353. Under PMF-driven conditions, proton transfer through DabB2 leads to the deprotonation or protonation of residues along the conduction pathway. These events possibly modulate the conformation of both the substrate tunnels and the catalytic site through the DabA2 transmembrane ‘finger-like’ motifs, thereby enabling efficient CO_2_ hydration and HCO₃⁻ export.

**Figure 6.**
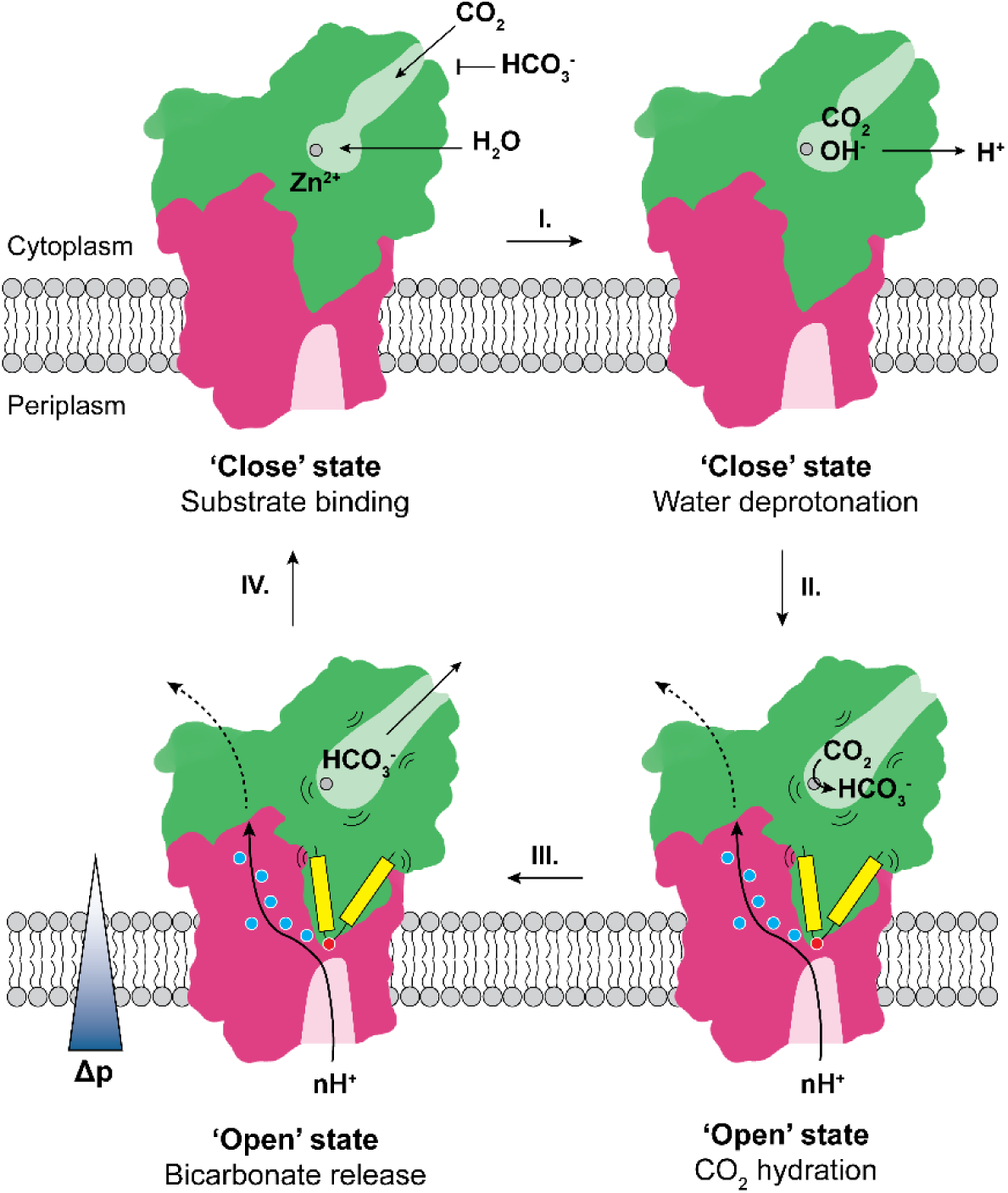
Hypothesized regulatory mechanism. In the ‘Close’ state, DabA2 binds CO_2_ and water molecules at the active site, meanwhile bicarbonate binding is sterically disfavored. The zinc-bound water deprotonates to form a hydroxide ion (Step **I.**) however, the active site likely remains catalytic inactive at this stage. The proton motive force (Δp) drives proton transfer across DabB2 via charged and polar residues (blue dots). (De-)Protonation of the DabA2 “finger-like” motifs (yellow) terminal residue Glu444 (red dot) and residues along the proton pathway possibly triggers structural rearrangement at the active and the substrate tunnel, hence activating CO_2_ hydration (Step **II.**) and subsequent release of bicarbonate formed (Step **III.)**. The complex returns to the ‘Close’ state, for substrate binding after releasing the protons and bicarbonate (Step **IV.**).

Our findings reveal a highly coordinated regulatory mechanism in which structural rigidity and dynamic tunnel gating converge to control catalysis in the DAC complex. PMF-dependent activation opens the substrate tunnels, enables active site reorganization and permits unidirectional product release. This establishes a structural basis for directionality, while also suppressing the thermodynamically favored reverse reaction of bicarbonate dehydration. This ensures that catalysis only proceeds under conditions of sufficient proton motive force, effectively linking carbon uptake to the energetic state of the cell. Unlike ferredoxin-dependent CO_2_ hydration in cyanobacterial NDH-1 complexes, DAB2 functions independently of electron transfer, representing a distinct adaptation for CO_2_ capture and conversion in chemolithoautotrophs inhabiting energy- and carbon-limited environments.

## Methods

### Cloning, overexpression and purification of DAB2

The genes *hneap_0212* and *hneap_0211* encoding the DAB2 complex were amplified from *Halothiobacillus neapolitanus* genomic DNA and cloned into pET-24d by Golden Gate Assembly with a 3C protease cleavage site and a twin-strep tag at the C-terminal of DabA2. For the construction of DAB2 fusion (*Dab2*), the stop codon of *hneap_0212* was removed and directly fused to the start codon of *hneap_0211* as one gene. After verified by sequencing, the plasmids were transformed into chemically competent *E. coli* BL21 (DE3). To generate variants of *Dab2*, specific point mutations were introduced by site-directed mutagenesis following standard protocol.

For protein overexpression, bacteria were cultivated at 37 °C in Luria-Bertani (LB) medium supplemented with 25 μg/mL kanamycin and 0.1 mM ZnSO4 until reaching OD600 0.7-0.8. The cultivation temperature was reduced to 16 °C and cells were harvested after 16 hours without addition of inducer to reduce protein aggregation.

Cell pellet was resuspended in lysis buffer (20 mM HEPES, 150 mM NaCl, 1 mM PMSF, pH 7.5), and lysed by passing through a microfluidizer (Microfluidics) at 12,000 PSI for three times. Cell debris was removed by centrifugation at 40,000 xg, 4 °C for 20 mins, and membrane vesicles were isolated from the supernatant by ultracentrifugation at 200,000 xg, 4 °C for 1 hour. The membrane pellet was used immediately or stored at - 80 °C until use. To purify DAB2, the pellet was homogenized in lysis buffer at 0.1-0.2 g wet weight per mL buffer. Protein was solubilized with 1% n-dodecyl-β-d-maltoside (DDM) and gently stirred at 4 °C for 1 hour. Insoluble material was removed by ultracentrifugation at 200,000 xg, 4 °C for 30 mins. The solubilized protein was incubated with pre-equilibrated Strep-Tactin resin (IBA Lifesciences) at 4 °C for 1 hour with gentle agitation. The resin was washed with 15 column volumes of wash buffer (20 mM HEPES, 150 mM NaCl, 0.03 % DDM, pH 7.5). The protein was eluted in wash buffer supplemented with 2.5 mM desthiobiotin and concentrated to 5-7 mg/mL using a 100 kDa cut-off Amicon centrifugal filter (Merck Millipore). The protein was further purified by size-exclusion chromatography (SEC) using a ÄKTA Pure system (Cytiva) and Superose 6 increase 10/200 GL column (Cytiva) in wash buffer. The column was calibrated using the HMW Gel Filtration Calibration Kits (Cytiva) according to the manufacture instruction. Peak fractions were collected and concentrated when necessary. The purity of the proteins was analyzed by SDS-PAGE. Protein was used immediately or supplemented with 10 % glycerol and stored at -80 °C. The same procedure was used to purify Dab2 and mutant variants.

### Nanodisc reconstitution

Membrane scaffold protein MSP1D1 was purified according to established protocol^45^. Chloroform dissolved *E. coli* polar lipid extract (Avanti Research) was evaporated under a gentle stream of nitrogen to from a thin lipid film. Residue organic solvent was removed under vacuum overnight. Dried lipid was hydrated in reconstitution buffer (20 mM HEPES, 150 mM NaCl, 0.1 M Na-cholate, pH 7.5) to 25 mg/mL and water bath sonicated (Diagenode) at high setting for 30 – 60 seconds to aid hydration. 32 uM of Dab2 was incubated with MSP1D1 and lipid at 1:2:40 molar ratio (assuming 800 g/mol for the lipid extract) at 4 °C with gentle agitation. The final reconstitution buffer contained 20 mM HEPES, 150 mM NaCl, 14 mM Na-cholate. A total of 0.25 g pre-equilibrated Bio-beads SM-2 resins (Bio-Rad) was added in two successions – at one hour and 3 hours of incubation. Afterward, the mixture was incubated for 2 more hours. Supernatant was collected and centrifuged at 14,000 xg for 10 mins at 4 °C to remove aggregates. The reconstituted protein was analyzed by Superose 6 increase 10/200 GL column in SEC buffer (20 mM HEPES, 150 mM NaCl, pH 7.5). Peak fractions were concentrated when necessary. Sample was immediately used to prepare cryo-EM grids.

### Cryo-EM sample preparation and data collection

For cryo-EM sample preparation, 4 µl of the protein sample (2 mg/mL) was applied to glow-discharged Quantifoil 2/1 Cu200 mesh grids, blotted for 6 s with force 6 in a Vitrobot Mark IV (Thermo Fisher) at 100% humidity and 4 °C, and plunge frozen in liquid ethane. For sample prepared under high CO_2_ condition (*Dab2-CO_2_*), saturated CO_2_ water (prepared by bubbling milli-Q water with solid CO_2_ at 4 °C for at least 30 minutes, until reaching pH 4.0) was added to the protein sample to final concentration of approximately 17 mM dissolved CO_2_ shortly before blotting. For sample prepared under high bicarbonate condition (*Dab2-HCO_3_^-^*), the protein was incubated in 100 mM NaHCO_3_ for 2 mins prior to blotting. Cryo-EM datasets were collected on a Titan Krios G4i electron microscope operated at 300 kV equipped with a Falcon 4i direct electron detector (Thermofisher scientific). Movies were collected at 165,000x magnification (0.73 Å pixel size) with a defocus range of -0.5 to -2.25 µm.

### Image processing

The processing pipeline for each dataset was summarized in supplementary Figures 2 to 4. All processing steps were carried out in cryoSPARC (v4.7.0)^46^. For *Dab2-ambient*, 24,092 EER movies were fractionated into 60 fractions per stack without upsampling and motion-corrected using Patch Motion Correction. Contrast transfer function (CTF) was calculated using Patch CTF Estimation. Roughly 200 initial particles were manually picked from a random subset of 100 corrected micrographs as templates to guide optimized blob picking using Blob Picker Tuner. ‘Junk’ particle picks were filtered using Inspect Particle Picks, while the remaining picks were extracted in a 360 pixels box, yielding ∼6 million particles and downsampled by 4 times, followed by 2D classification. Desirable 2D classes (∼3 million particles) were selected for ab-initio reconstruction in 2 classes. The class containing good particles were refined by Non-uniform Refinement using particles extracted in full size and 3D classified into 6 classes to further resolve poor particles and heterogeneity. The class with the most prominent CO_2_-like densities and zinc-bound water was subjected to Heterogenous Refinement in 2 classes. The ‘best’ class was further refined by Non-uniform Refinement and Local Refinement with a mask on the entire protein, excluding the nanodisc density, after particles were polished using Reference-based Motion correction. This resulted in the final reconstruction of 2.64 Å (global FSC = 0.143) with 254,703 particles. A similar strategy was utilized to process the *Dab2-CO_2_* and *Dab2-HCO_3_^-^* datasets.

For *Dab2-CO_2_*, ∼3 million initial particles were extracted from 20,458 micrographs after motion correction, CTF estimation and excluding movies with CTF fits worse than 3.5 Å. Approximately 1.6 million particles were selected after 2D classification to generate 2 classes of an-initio densities. The ‘good’ class was passed to 3D classification in 5 classes and heterogenous refinement in 2 classes. Particles from the ‘best’ class was polished and used for Non-uniform Refinement followed by Local Refinement, yielding the final reconstruction of 2.72 Å (global FSC = 0.143) with 231,139 particles.

For Dab2-HCO_3_ ^-^, ∼3 million initial particles were extracted from 13,704 micrographs after motion correction, CTF estimation and excluding movies with CTF fits worse than 3.5 Å. After 2D classification, ∼2 million particles were selected to generate 2 classes of an-initio densities. The ‘good’ class was 3D classified into 5 classes. The class that contained density corresponding to a zinc-bound bicarbonate was subjected to a round of Non-uniform Refinement. Preferred orientation was mitigated by removing over-populated particles using the Rebalance Orientations tool. The complex was locally refined with a mask on the entire protein using the normalized particle sets, resulting in the final reconstruction of 3.22 Å (global FSC = 0.143) with 226,711 particles.

### Model building and refinement

An AlphaFold2 (v1.5.5)^47^ predicted model of the DAB2 complex was rigid-body fitted into the densities in ChimeraX (v1.10)^48^ and subjected to a round of real space refinement in Phenix (v1.20.1)^49^ to create an initial model. Ligands were modelled manually in COOT (v0.9.8)^50^ and the initial model was refined with several rounds of Phenix real space refinement and manual refinement. Water molecules were added using douse, followed by manual inspections and refinement. The statistics of all cryo-EM data acquisition and refinement are summarized in Supplementary Table 1.

### Site-directed mutagenesis and complementation assay

To investigate the importance of specific amino acid residues, *hneap_0212* and *hneap_0211* were cloned into pBAD30 which offers a tight genetic regulation for complementation experiments. The plasmid was engineered with kanamycin resistance to enhance antibiotic stability. Mutations were introduced by site-directed mutagenesis using Golden Gate Assembly^51^ and the sequenced plasmids were transformed into *E. coli* Lemo21 lacking carbonic anhydrases (Δ*can*Δ*cynT*)^14^. This strain could only grow under ambient air if complemented by carbonic anhydrases. Bacteria were cultivated overnight in LB medium supplemented with 25 μg/mL kanamycin under 5 % CO_2_ atmosphere, then inoculated into fresh LB medium with antibiotic and 0.1% L-arabinose. Growth (absorbance at 600 nm) was monitored using a microplate reader (Tecan) at 37 °C, under ambient atmosphere with agitation. Growth yield was measured at the 10-hour time point when the wild-type strain entered stationary phase and the results were presented relative to that of the wild-type.

### Sodium transporter activity assay

Sodium transporter activity was evaluated based on the previously reported method^44^. Genes coding the key sodium transporters (*nhaA* and *nhaB*) were deleted by scarless allelic exchange in *E. coli* MG1655^52^. In brief, 1000 bp flanking sequences upstream and downstream of *nhaA* and *nhaB* were cloned into the suicide plasmid pKOV and transformed into *E. coli* MG1655. Successive homologous recombinations were performed at 30 °C, followed by plasmid curing at 43 °C and sucrose counter selection. Gene deletion was verified by colony PCR. The double mutant strain was transformed with pBAD30 expressing wild-type DAB2 and routinely cultivated in LB-K (LB with NaCl replaced by KCl). To test the sodium transporter activity, overnight culture was inoculated in LB medium containing 0.1 M NaCl, 25 μg/mL kanamycin and 0.1% L-arabinose. Bacterial growth was monitored as described above.

To verify DAB2 sodium dependency, Δ*can*Δ*cynT E. coli* expressing wild-type DAB2 was cultivated in M9 minimal medium^53^ prepared with sodium salts (M9-Na) or potassium salts (M9-K), supplemented with 0.4% glycerol. Bacteria were cultivated overnight in M9-K supplemented with 25 μg/mL kanamycin, under 5 % CO_2_ to deplete cellular sodium prior to inoculation into fresh M9-Na or M9-K. Sodium dependency, as a function of growth was monitored as described above.

### Carbonic anhydrase activity assay

Carbonic anhydrase activity was measured based on the Wilbur-Anderson method^54^ using a UV/Vis spectrophotometer (JASCO V-750) at room temperature. 0.3 mL Ice-cold CO_2_ saturated water was mixed with 0.7 mL assay buffer to initiate CO_2_ hydration. The reaction mixture contained 5 nM Bovine carbonic anhydrase II (Sigma) or 500 nM DDM purified *Dab2*, 20 mM Tris, 150 mM NaCl, 0.03% DDM, 100 µM phenol red, pH 8.3. Acidification as a result of CO_2_ hydration was monitored at 558 nm absorbance and 0.2 second time-resolution.

### Fourier transform infrared spectroscopy

All experiments were performed on hydrated protein films in attenuated total reflection (ATR) configuration using a FTIR spectrometer (Bruker Tensor27) equipped with a mercury cadmium telluride (MCT) detector cooled by liquid N_2_. All data were recorded with a spectral resolution of 2 cm^-1^ at 80 kHz scanning velocity. For 25 co-additions of interferometer scans in forward/backward direction, a temporal resolution of 5 s was achieved. The hydration reaction was started by changing the atmosphere from 100% N_2_ to 90% N_2_ and 10% CO_2_ (1 L/min), as reported earlier^29^. For each experiment, 1µL of 100–200 µM protein solution (ECCA, BSA, Dab2) was used and comparable hydration levels were adjusted (Fig. S13) Moreover, all experiments were conducted under ambient temperature and pressure, and in the dark. For the kinetic evaluation, FTIR difference spectra between 1800–1200 cm^-1^ were fitted with contributions from HCO_3_^−^ at 1614, 1360, and 1302 cm^-1^ with a FWHM of 53, 48, and 64 cm^-1^, respectively. This allowed calculating the HCO_3_^−^ peak area, which is plotted against time in Fig. 3c as a measure of catalytic activity. Amide band changes at 1650 and 1545 cm^-1^ (FWHM 60 and 55 cm^-1^) are considered in the fit (Fig. S13); this unspecific behavior does not affect the analysis of HCO_3_^−^ kinetics. We used home-written software for fitting, as reported earlier^29^.

### Inductively coupled plasma mass spectrometry

For metal ion determination, inductively coupled plasma-triple quadrupole mass spectrometry (ICP-QQQ-MS) was performed. Briefly, purified and desalted protein samples were subjected to acid digestion by incubating them in 11% (v/v) HNO₃ (Suprapur grade) for 3 hours at 80°C. After total hydrolysis, the samples were diluted with ultrapure water to achieve a final HNO₃ concentration of 2% (v/v). Calibration standards, ranging from 0.005 µg/L to 500 µg/L, were prepared by serially diluting the ICP multi-element standard solution Merck XVI (Merck Millipore) in 2% (v/v) HNO₃. To ensure accuracy, a rhodium internal standard was added to all samples, resulting in a final concentration of 1 µg/L. Metal analysis was conducted using a high-resolution ICP-QQQ-MS system (Agilent 8800, Agilent Technologies) in direct infusion mode with an integrated auto-sampler. The injection system included a Peltier-cooled (2°C) Scott-type spray chamber equipped with a perfluoroalkoxy alkane (PFA) nebulizer, operating at 0.3 revolutions per second (rps) for 45 seconds with an internal tube diameter of 1.02 mm. Multiple metals were quantified simultaneously using the Merck XVI standard solution. To reduce polyatomic interferences, the Octopole Reaction System (ORS3) with a collision/reaction cell (CRC) was utilized. Helium (2.5 mL/min) and hydrogen (0.5 mL/min) were introduced into the CRC as collision/reaction gases, while argon was used as the carrier gas at a flow rate of 2.7 mL/min. For each metal, the first (Q1) and second (Q2) quadrupoles were set to the same m/z value, with an integration time of 1 second under auto-detector mode. All measurements were conducted in technical triplicates and normalized using the internal standard, and additional parameters were optimized via the auto-tune function in the MassHunter 4.2 software (Agilent Technologies).

### Sequence conservation analyses

Amino acid conservation analysis was performed using the ConSurf server^55^. Protein sequences of DabA2 and DabB2 were searched against the NCBI non-redundant (nr) protein database, in which 150 sequences (for DabA2) and 100 sequences (for DabB2) were sampled from the list of homologues for the calculation. Structural-based sequence alignment was carried out using the DALI server^20^ and visualized in Jalview^56^.

### Total proteome analysis

Bacteria were cultivated as described in the complementation assay above. The cell pellets were resuspended in 300 μl lysis buffer (2% sodium lauroyl sarcosinate (SLS), 100 mM ammonium bicarbonate) and heated at 90°C for 40 min. The protein amount was determined by bicinchoninic acid protein assay (Thermo Scientific). Proteins were incubated with 5 mM Tris(2-carboxyethyl) phosphine (Thermo Fischer Scientific) and 10 mM Chloroacetamide at 90°C for 15 min (Sigma Aldrich) and further processed with SP3 (ref.^57^). Proteins were bound to 4 µl SP3 beads (40% v/v bead stock) in presence of 70 % acetonitrile for 15 min at room temperature, followed by two washes of beads with 70 % ethanol and an additional wash with acetonitrile. After removal of the supernatant, 1 µg trypsin in 100 mM NH_4_HCO_3_ was added to the beads and digested shaking overnight at 30°C. Digested proteins were harvested and purified using C18 solid phase extraction. Peptides were finally dried, reconstituted in 0.1 % trifluoroacetic acid (TFA) and analyzed using liquid-chromatography-mass spectrometry carried out on an Exploris 480 instrument connected to an VanquishNeo and a nanospray flex ion source (all Thermo Scientific). Peptide separation was performed on a reverse phase HPLC column (75 μm x 26 cm) packed in-house with C18 resin (1.9 μm Reprosil-AQ; Dr. Maisch). The following separating gradient was used: 100% solvent A (0.1% formic acid) to 40% solvent B (99.85% acetonitrile, 0.15% formic acid) over 32 minutes at a flow rate of 300 nl/min. The direct injection setup was applied.

MS raw data was acquired in data independent acquisition (DIA) mode. The funnel RF level was set to 40. Full MS resolution was set to 120.000 at m/z 200. AGC target value for fragment spectra was set at 3000%. 45 windows of 14 Da were used with an overlap of 1 Da between m/z 320-950. Resolution was set to 15,000 and IT to 22 ms. Stepped HCD collision energy of 25, 27.5, 30 % was used. MS1 data was acquired in profile, MS2 DIA data in centroid mode.

Analysis of DIA data was performed using the DIA-NN version 1.9 (ref.^58^) using a uniprot protein database from *E.coli* BL21 including target proteins to generate a data set specific spectral library for the DIA analysis. The neural network based DIA-NN suite performed noise interference correction (mass correction, RT prediction and precursor/fragment co-elution correlation) and peptide precursor signal extraction of the DIA-NN raw data. The following parameters were used: Full tryptic digest was allowed with two missed cleavage sites, and oxidized methionines (variable) and carbamidomethylated cysteins (fixed). Match between runs and remove likely interferences were enabled. The precursor FDR was set to 1%. The neural network classifier was set to the single-pass mode, and protein inference was based on genes. Quantification strategy was set to any LC (high accuracy). Cross-run normalization was set to RT-dependent. Library generation was set to smart profiling. DIA-NN outputs were further evaluated using the SafeQuant script^59,60^ modified to process DIA-NN outputs

## Supplementary tables and figures

**Supplementary Table 1.** Cryo-EM data collection, refinement and validation statistics.

|  | Dab2-Ambient<br>(EMDB-53925)<br>(PDB 9RD0) | Dab2-CO<sub>2</sub><br>(EMDB-53930)<br>(PDB 9RD9) | Dab2-HCO<sub>3</sub><sup>-</sup><br>(EMDB-53929)<br>(PDB 9RD8) |
| --- | --- | --- | --- |
| Data collection and processing |  |  |  |
| Magnification | 165,000X | 165,000X | 165,000X |
| Voltage (kV) | 300 | 300 | 300 |
| Electron exposure (e<sup>-</sup>/Å<sup>2</sup>) | 55 | 55 | 55 |
| Defocus range (μm) | -0.5 to -2.25 | -0.5 to -2.25 | -0.5 to -2.25 |
| Pixel size (Å) | 0.73 | 0.73 | 0.73 |
| Symmetry imposed | C1 | C1 | C1 |
| Initial particle images (no.) | 6,304,235 | 2,863,711 | 3,262,632 |
| Final particle images (no.) | 254,703 | 231,193 | 226,711 |
| Map resolution (Å) | 2.64 | 2.72 | 3.22 |
| FSC threshold | 0.143 | 0.143 | 0.143 |
| Map resolution range (Å) | 2.3-3.0 | 2.4-3.1 | 2.8-4.0 |
| Refinement |  |  |  |
| Model resolution (Å) | 2.6 | 2.7 | 3.2 |
| FSC threshold | 0.143 | 0.143 | 0.143 |
| Map sharpening *B* factor (Å<sup>2</sup>) | -120.1 | -115.1 | -90.9 |
| Model composition |  |  |  |
| Non-hydrogen atoms | 10,687 | 10,539 | 10,342 |
| Protein residues | 1,320 | 1,317 | 1,312 |
| Ligands | 10 | 10 | 7 |
| *B* factors (Å<sup>2</sup>) |  |  |  |
| Protein (mean) | 29.82 | 45.88 | 17.24 |
| Ligand (mean) | 36.46 | 53.51 | 35.48 |
| R.m.s. deviations |  |  |  |
| Bond lengths (Å) | 0.002 | 0.003 | 0.005 |
| Bond angles (°) | 0.456 | 0.514 | 0.963 |
| Validation |  |  |  |
| MolProbity score | 1.35 | 1.38 | 1.59 |
| Clashscore | 5.00 | 6.11 | 6.62 |
| Poor rotamers (%) | 0.28 | 0.38 | 0.29 |
| Ramachandran plot |  |  |  |
| Favored (%) | 97.56 | 97.78 | 96.54 |
| Allowed (%) | 2.37 | 2.14 | 3.30 |
| Disallowed (%) | 0.08 | 0.08 | 0.15 |

**Supplementary Table 2.** Polar interactions between DabA2 and DabB2 identified by the PISA server.

**Hydrogen bond**
| **DabB2** | **Distance (Å)** | **DabA2** | **DabB2** | **Distance (Å)** | **DabA2** |
| --- | --- | --- | --- | --- | --- |
| Tyr109 | 3.44 | Trp649 | Ser356 | 3.17 | Asn530 |
| Tyr109 | 3.35 | Asn530 | Thr358 | 3.39 | Gly738 |
| Gln112 | 3.17 | Gln651 | Asp359 | 2.72 | Arg534 |
| Asp115 | 2.71 | Arg653 | His360 | 2.66 | Ser464 |
| Arg167 | 3.06 | Glu744 | Arg362 | 2.80 | Gln545 |
| Gln247 | 3.27 | Asn746 | Arg362 | 2.81 | Arg387 |
| Met249 | 2.98 | Asn746 | Gln363 | 3.28 | Arg387 |
| Glu250 | 3.01 | Asn746 | Ala364 | 2.55 | Gln423 |
| Thr253 | 2.93 | VAL731 | Lys365 | 2.96 | Glu390 |
| Leu304 | 3.17 | Leu154 | Lys368 | 3.44 | Gly389 |
| Leu304 | 2.85 | TRP155 | Glu431 | 2.82 | His420 |
| Ser307 | 2.92 | Ala145 | Gln490 | 2.62 | Ser435 |
| Ser307 | 2.97 | Gly737 | Gln490 | 3.23 | Ala436 |
| Ser307 | 3.17 | Asp149 | Gln490 | 2.78 | Ala437 |
| Ile309 | 3.46 | Asn734 | Asn510 | 3.37 | Asn146 |
| Lys310 | 3.04 | Asn734 | Asn510 | 2.64 | Gln142 |
| Gln320 | 2.95 | CYS439 | Asn511 | 3.48 | Ser178 |
| His349 | 2.78 | Asp457 | Asn511 | 3.05 | Gln142 |
| Leu352 | 3.26 | Asn530 | Asn511 | 3.53 | Asn176 |
| Gly355 | 3.41 | Asn530 | Arg521 | 3.16 | Ala181 |

**Salt bridge**
| **DabB2** | **Distance (Å)** | **DabA2** | **DabB2** | **Distance (Å)** | **DabA2** |
| --- | --- | --- | --- | --- | --- |
| Arg169 | 3.79 | Glu134 | Asp359 | 3.24 | Arg534 |
| Arg169 | 3.71 | Glu138 | Asp359 | 2.72 | Arg534 |
| His349 | 2.78 | Asp457 | Asp359 | 2.82 | Arg534 |
| Arg167 | 3.57 | Glu744 | Glu431 | 3.47 | His420 |
| Arg167 | 3.88 | Glu744 | Glu431 | 2.82 | His420 |
| Arg167 | 3.62 | Glu744 | Glu500 | 3.91 | Arg171 |

**Supplementary Figure 1.**
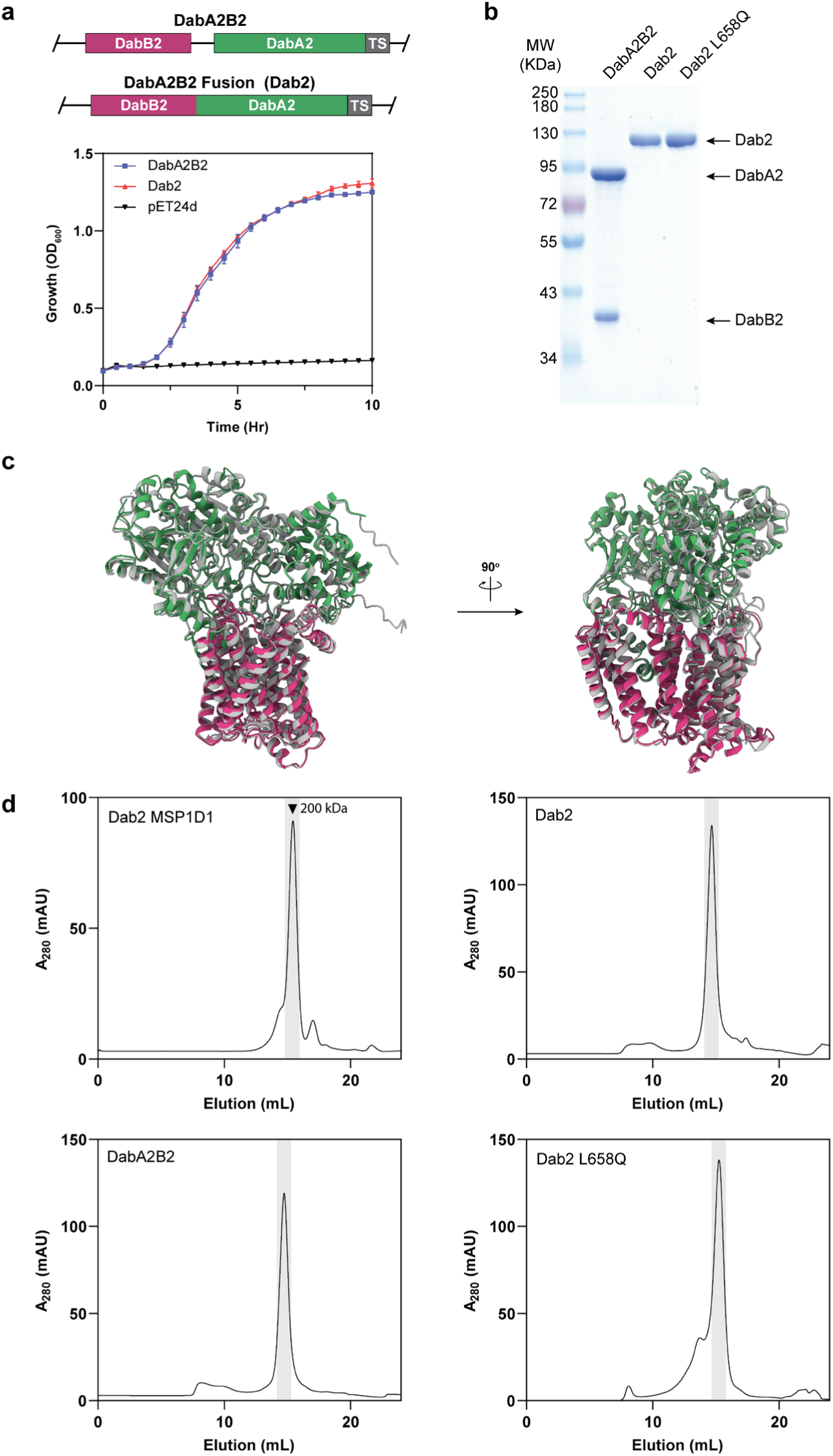
Construction and purification of Dab2. **a** Construct designs for purification of the wild-type DabA2B2 complex and the fusion variant (Dab2; top). TS: twin-strep tag. Both DabA2B2 and Dab2 were able to complement CA deficient *Escherichia coli* under low CO_2_ condition (0.04%) to a similar extend (bottom). Strain transformed with empty plasmid pET24d was used as a negative control. Data points and error bars represent means and standard deviations, respectively (n = 4 biological replicates). **b** SDS-PAGE of purified proteins after size exclusion chromatography. **c** Superposition of AlphaFold predicted model of DabA2B2 (gray) on the Dab2 fusion protein (colored as in Fig. 1; RMSD = 1.17 Å). **d** Size-exclusion chromatograms of the purified proteins in 0.03% DDM and after nanodisc (MSP1D1) reconstitution. Fractions under the gray area were collected.

**Supplementary Figure 2.**
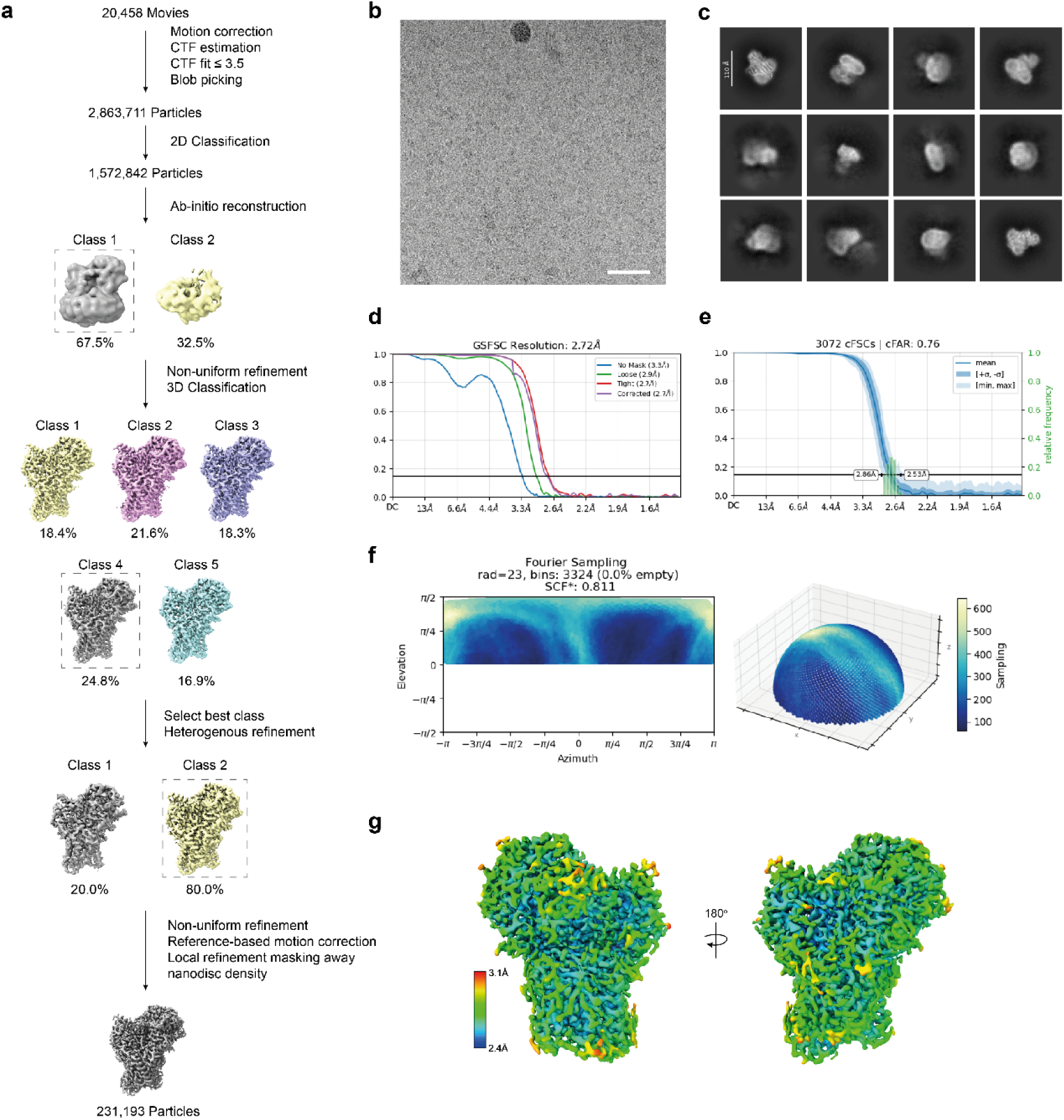
Cryo-EM processing workflow for *Dab2-CO_2_*. **a** Overview of the data processing pipeline (see methods). **b** Representative micrograph acquired on Falcon 4i Detector (scale bar, 50 nm). 20,458 micrographs were collected. **c** Representative 2D class averages. **d** Fourier shell correlation (FSC) curves before and after applying masks. Global resolution is reported at FSC = 0.143. **e** Directional resolution and **f** Sampling Compensation Factor (SCF*) of the final reconstruction. **g** Local resolution (Å) presented in front and back views.

**Supplementary Figure 3.**
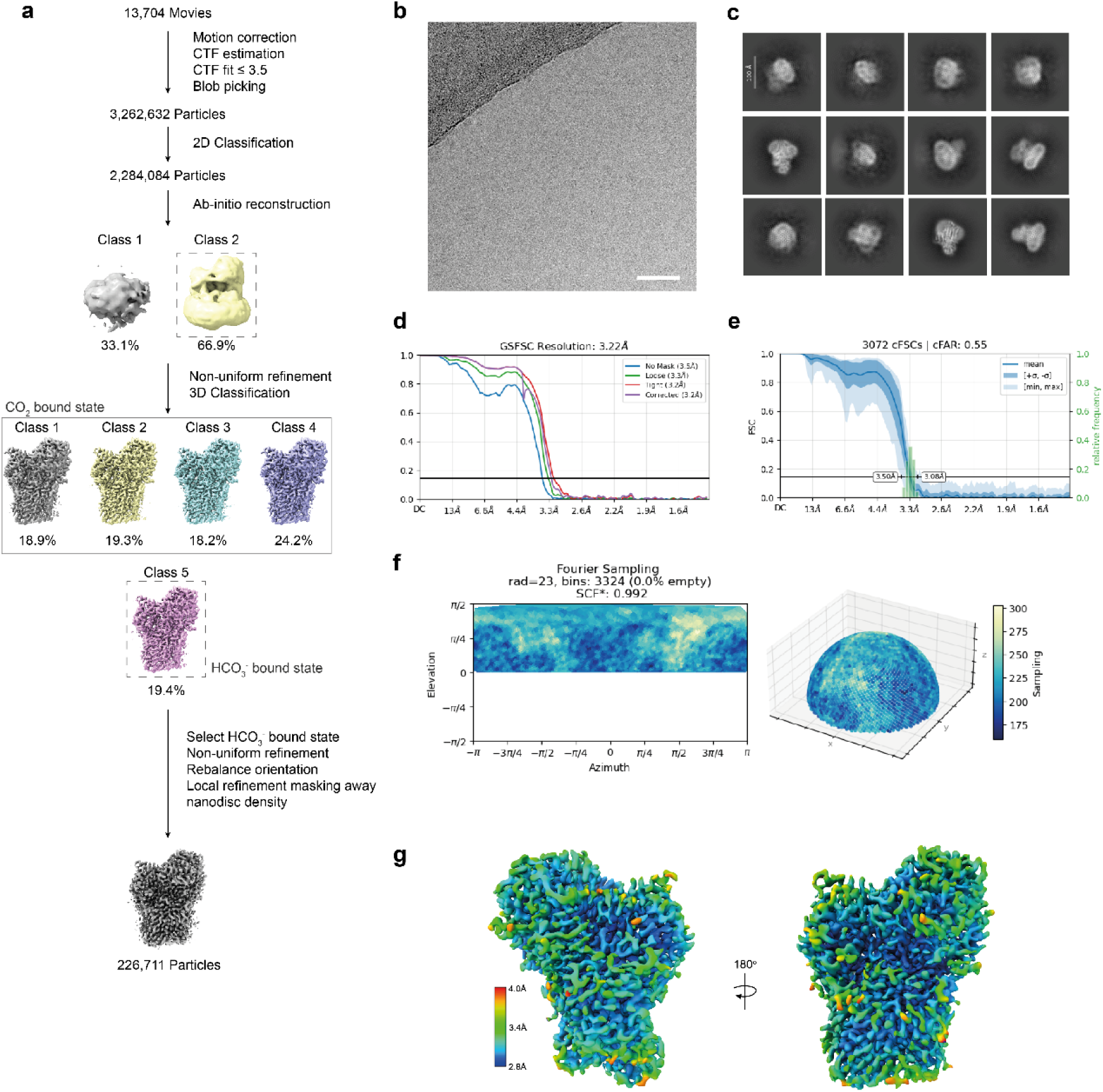
Cryo-EM processing workflow for *Dab2-HCO_3_^-^*. **a** Overview of the data processing pipeline (see methods). **b** Representative micrograph acquired on Falcon 4i Detector (scale bar, 50 nm). 13,704 micrographs were collected. **c** Representative 2D class averages. **d** Fourier shell correlation (FSC) curves before and after applying masks. Global resolution is reported at FSC = 0.143. **e** Directional resolution and **f** Sampling Compensation Factor (SCF*) of the final reconstruction. **g** Local resolution (Å) presented in front and back views.

**Supplementary Figure 4.**
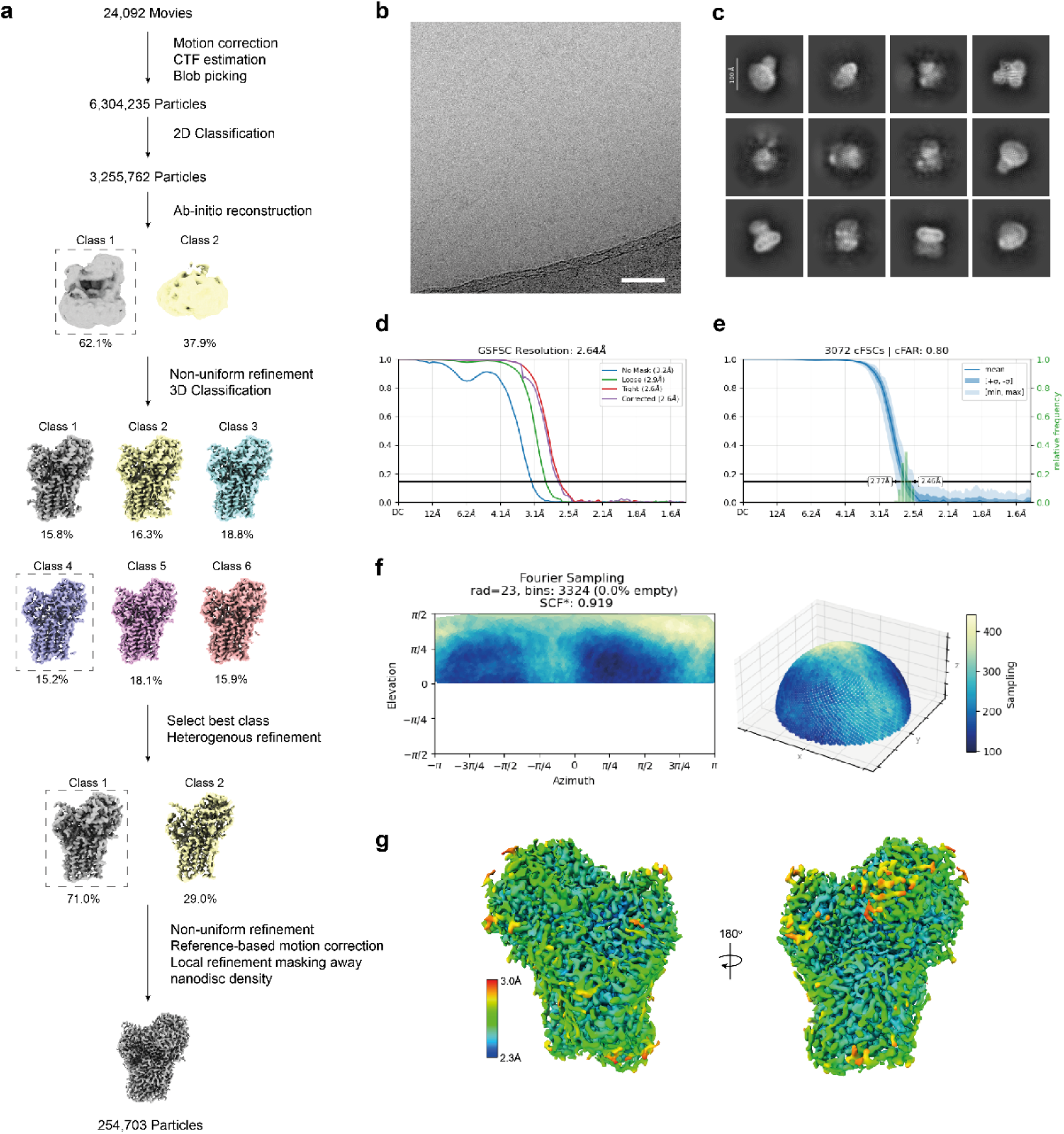
Cryo-EM processing workflow for Dab2-Ambient. **a** Overview of the data processing pipeline (see methods). **b** Representative micrograph acquired on Falcon 4i Detector (scale bar, 50 nm). 24,092 micrographs were collected. **c** Representative 2D class averages. **d** Fourier shell correlation (FSC) curves before and after applying masks. Global resolution is reported at FSC = 0.143. **e** Directional resolution and **f** Sampling Compensation Factor (SCF*) of the final reconstruction. **g** Local resolution (Å) presented in front and back views.

**Supplementary Figure 5.**
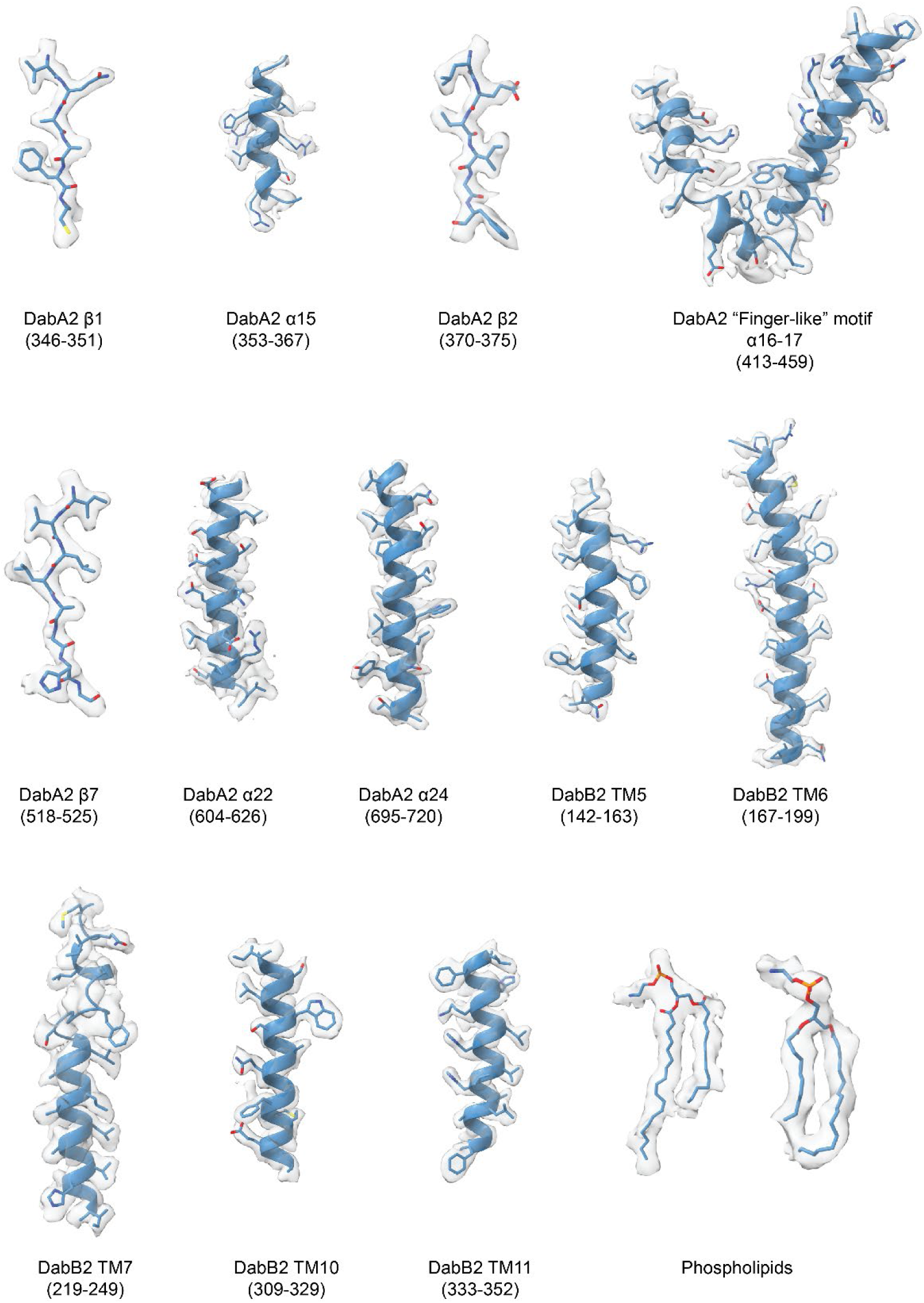
Representative density fitting. Density fitting of representative structures in DabA2, DabB2 and phospholipids.

**Supplementary Figure 6.**
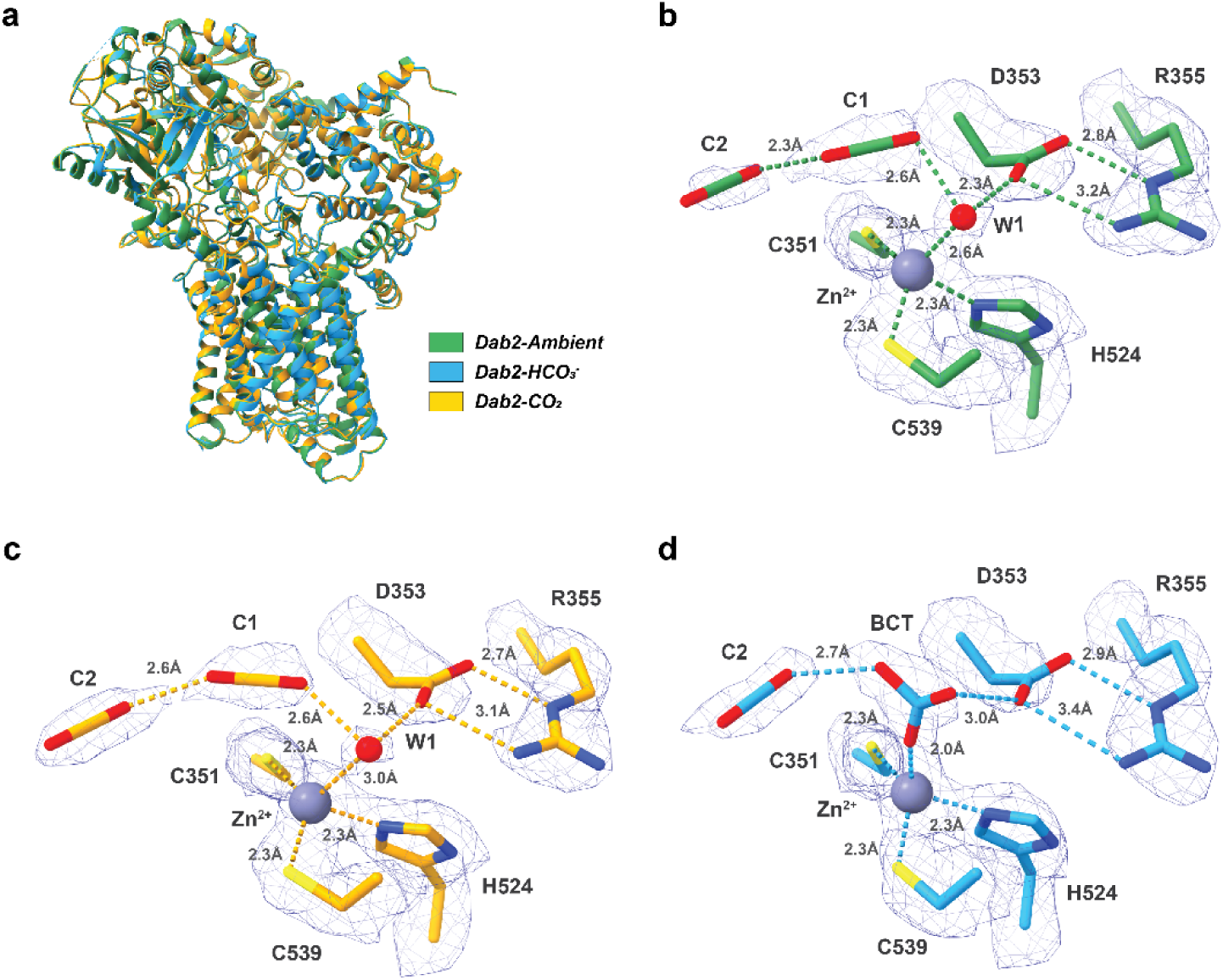
Comparison of *Dab2* under different conditions. **a** Superposition of *Dab2-ambient* (green), *Dab2-HCO_3_^-^* (blue), and *Dab2-CO_2_* (yellow). **b**-**d** Active site architecture of *Dab2* under different conditions, colored as in **a** and the density fitting. Maps are displayed at a comparable threshold (*Dab2-Ambient*: 7 σ; *Dab2-CO_2_*: 9.2 σ; *Dab2-HCO_3_^-^*: 6.5 σ). Dashes illustrate distance between interacting atoms and ions.

**Supplementary Figure 7.**
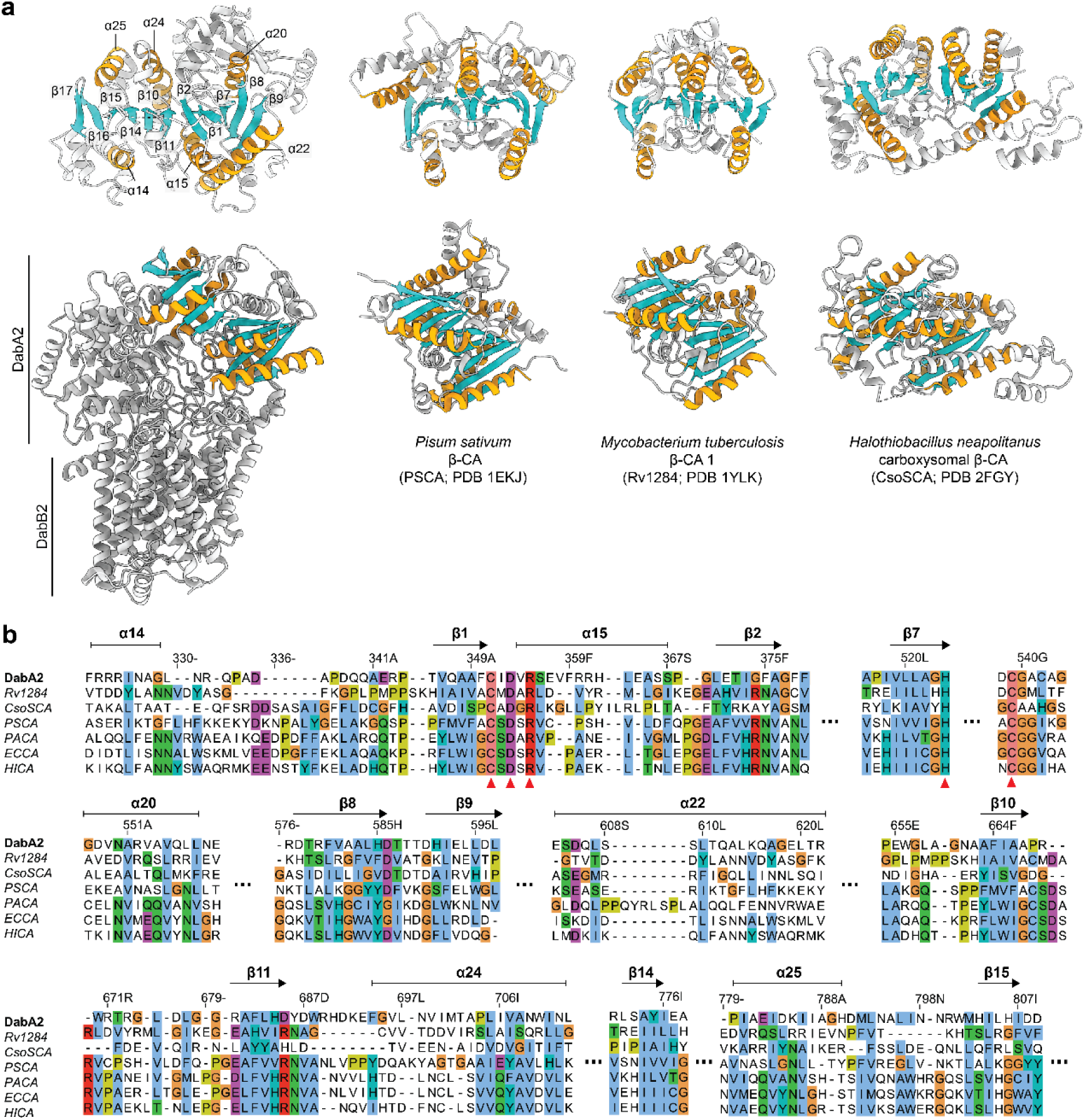
DabA2 catalytic domain structurally mimicked β-CAs. **a** Structural comparison between *Dab2* and representative β-CAs, shown in top and side views. DabA2 catalytic domain was characterized by two β-CA-like Rossmann folds, colored by secondary structure (orange: α-helix; cyan: β-sheet). The first fold consisted of four parallel β-sheets in the order of 2-1-3-4 (β2-β1-β7-β8) and an antiparallel sheet β9. β3-β6 formed the scaffold for the “finger-like” motifs (Supplementary Fig. 6). The second fold had an additional anti-parallel β-sheets, arranged in the ordered of β11-β10-β14-β15-β16-β17. β12 and β13 formed part of the interacting interface with DabB2. **b** Structure-based sequence alignment of DabA2 and β-CAs. Rv1284: *Mycobacterium tuberculosis* β-CA (PDB 1YLK); CsoSCA: *Halothiobacillus neapolitanus* β-CA (PDB 2FGY); PSCA: *Pisum sativum* β-CA (PDB 1EKJ); PACA: *Pseudomonas aeruginosa* β-CA (PDB 5BQ1); ECCA: *E. coli* β-CA (PDB 2ESF) HICA: *Haemophilus influenzae* β-CA (PDB 2A8D). Alignment is visualized in Jalview, and colored by the *Clustal* coloring scheme. Strictly conserved residues are marked by red triangles. DabA2 residues number and secondary structure are shown above the alignment. Un-aligned regions are trimmed and represented by dotted lines.

**Supplementary Figure 8.**
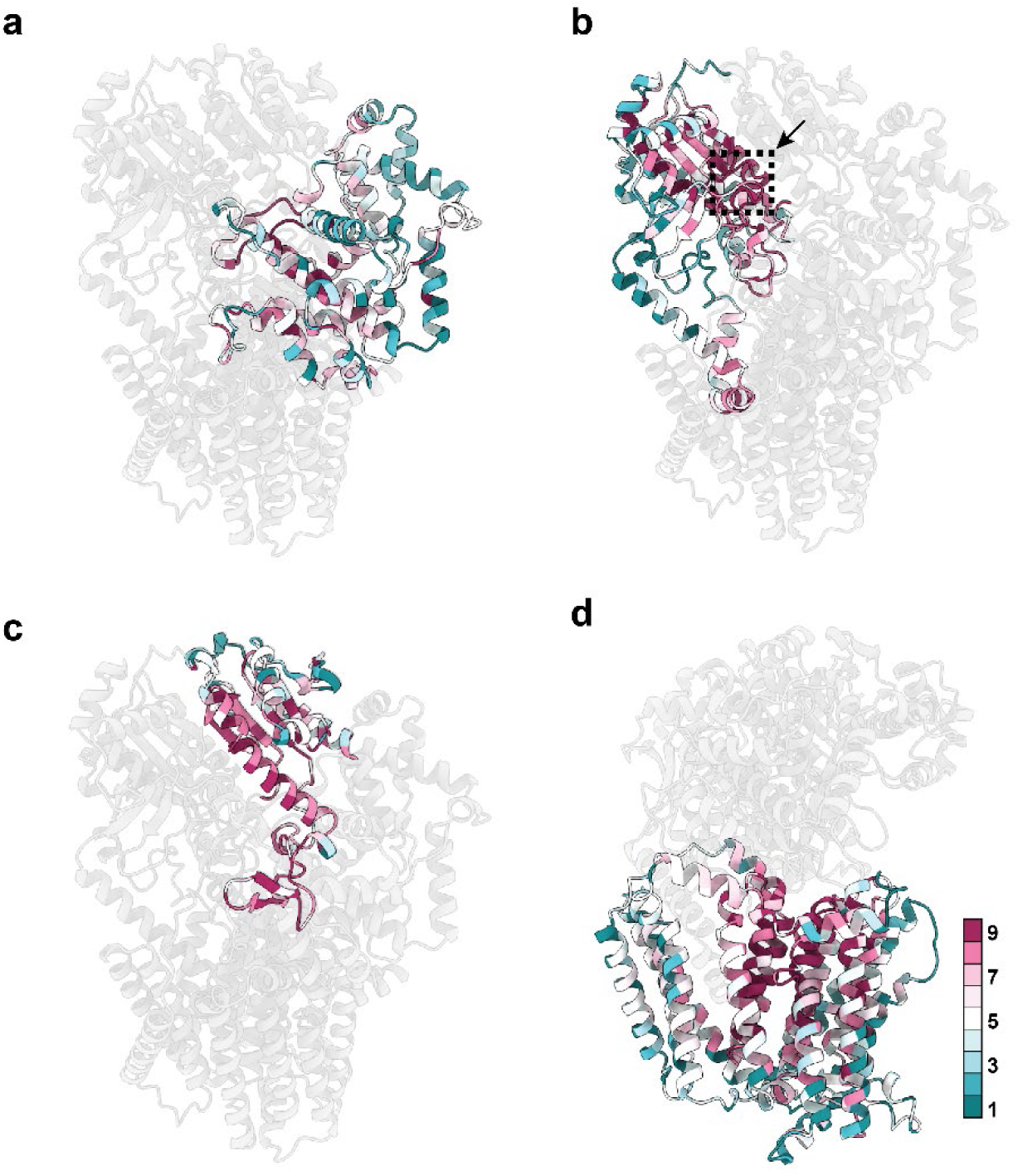
Amino acid conservation in DAB2. Sequence conservation of **a** N-terminal domain, **b** β-CA-like I domain, **c** β-CA-like II domain of DabA2 and **d** DabB2 calculated using the ConSurf server. Residues are colored by conservation level from highest (9) to lowest (1). **b** Arrow indicates position of the active site.

**Supplementary Figure 9.**
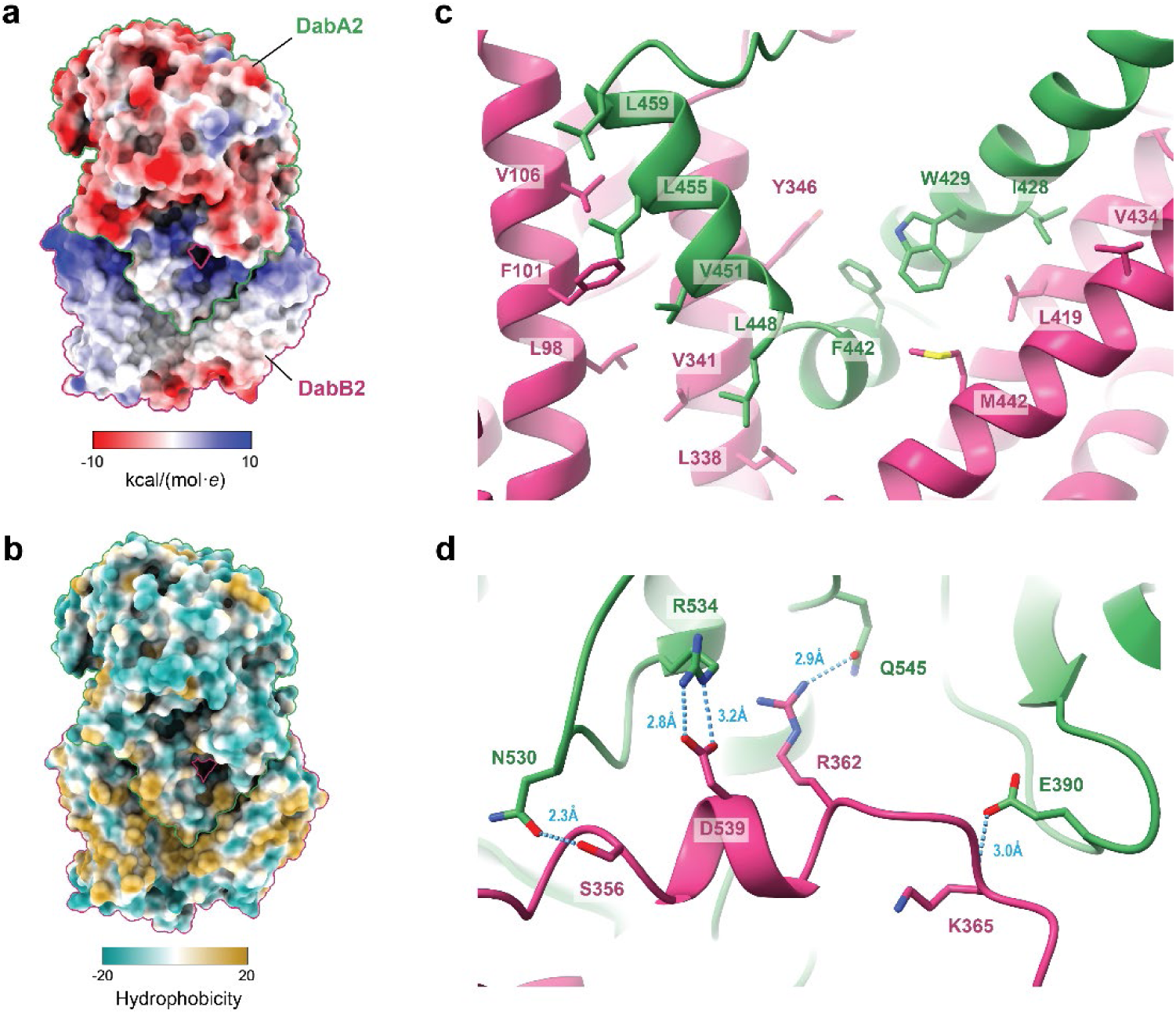
Interactions between DabA2 and DabB2. **a** Surface coulombic electrostatic potential and **b** hydrophobicity calculated by ChimeraX. Binding of DabA2 with DabB2 involved interactions at the cytoplasmic interface and hydrophobic interactions between DabB2 transmembrane helixes and DabA2 “finger-like” motif. **c** Overview of side chain hydrophobic interactions between DabB2 and DabA2 “finger-like” motif. **d** Representative hydrogen-bonds and salt bridge between DabA2 and DabB2 cytoplasmic helix (α2). See Supplementary Table 2 for the list of polar interactions.

**Supplementary Figure 10.**
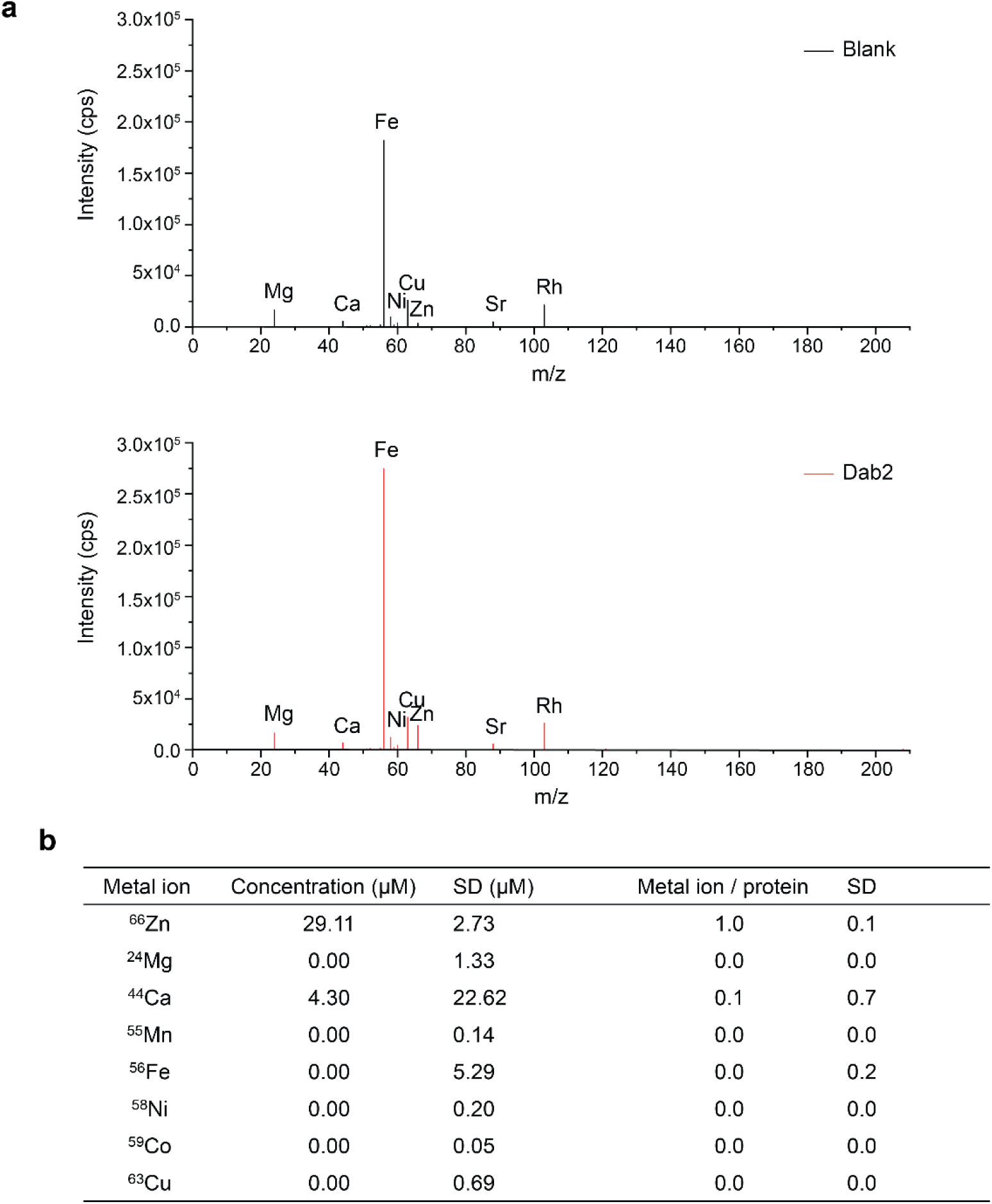
Inductively coupled plasma mass spectrometry. **a** Representative ICP-MS spectra of Dab2 and blank. **b** Concentration of metal ion detected from 30.6 µM protein and the corresponding metal-to-protein ratio. Means and standard deviations of three technical replicates are reported.

**Supplementary Figure 11.**
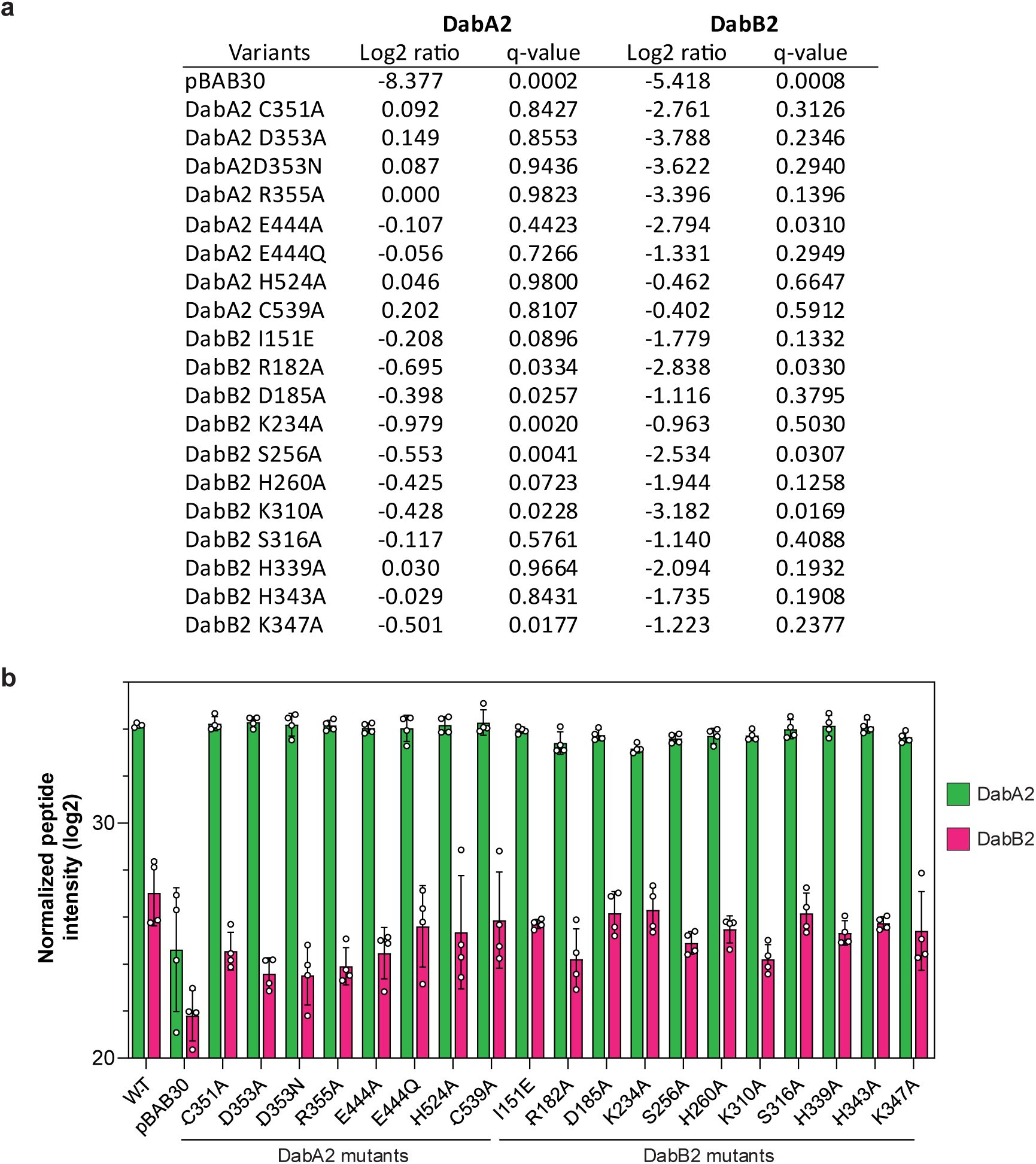
Protein expression level of Dab2 variants determined by LC-MS. **a** Log2 ratio of the normalized and imputed mean of Dab2 variants peptide intensity to the wild-type, expressed from the pBAD30 plasmid. See Materials and Methods for details. Lysate from strain carrying the empty pBAD30 plasmid was utilized as a ‘non-expressing’ negative control. Log2 ratio of zero indicates identical expression level as the wild-type complex. **b** Sum of peptide intensity for each variant imputed and normalized to the median protein sum of the total intensity distribution. The values were used to obtain panel **a**. Bar heights and error bars represent means and standard deviations, respectively (n = 4 biological replicates).

**Supplementary Figure 12.**
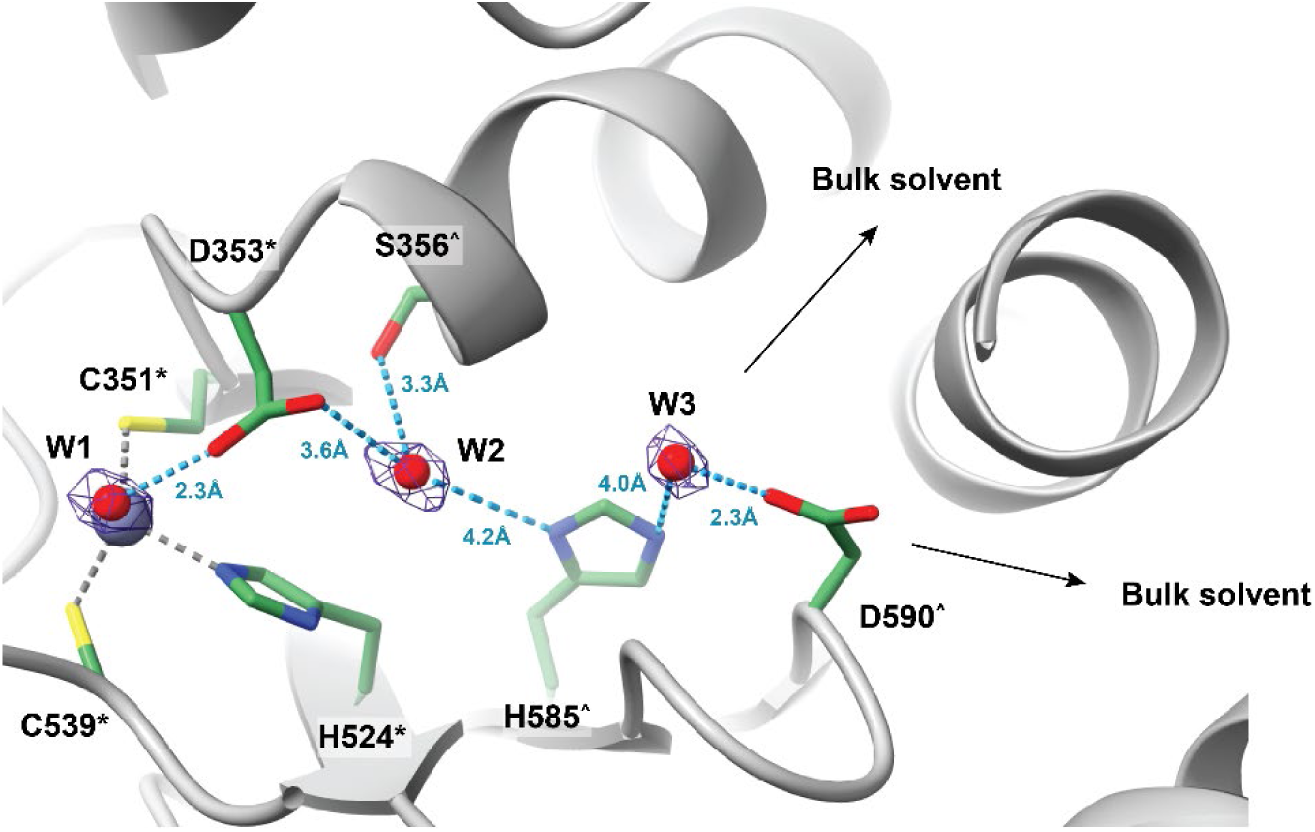
Active site water network. CO_2_ hydration is initiated by deprotonation of a zinc-bound water into a hydroxide ion. The water network (blue dashes) and charged residues connecting the active site and the bulk solvent may serve as a putative pathway for removing the proton. Densities of water molecules are displayed at 7 σ. Key residues are shown in green. Residues strictly conserved or conserved at ≥ 90% sequence identity among DabA2 homologues are marked by asterisk (*) and circumflex (^) respectively.

**Supplementary Figure 13.**
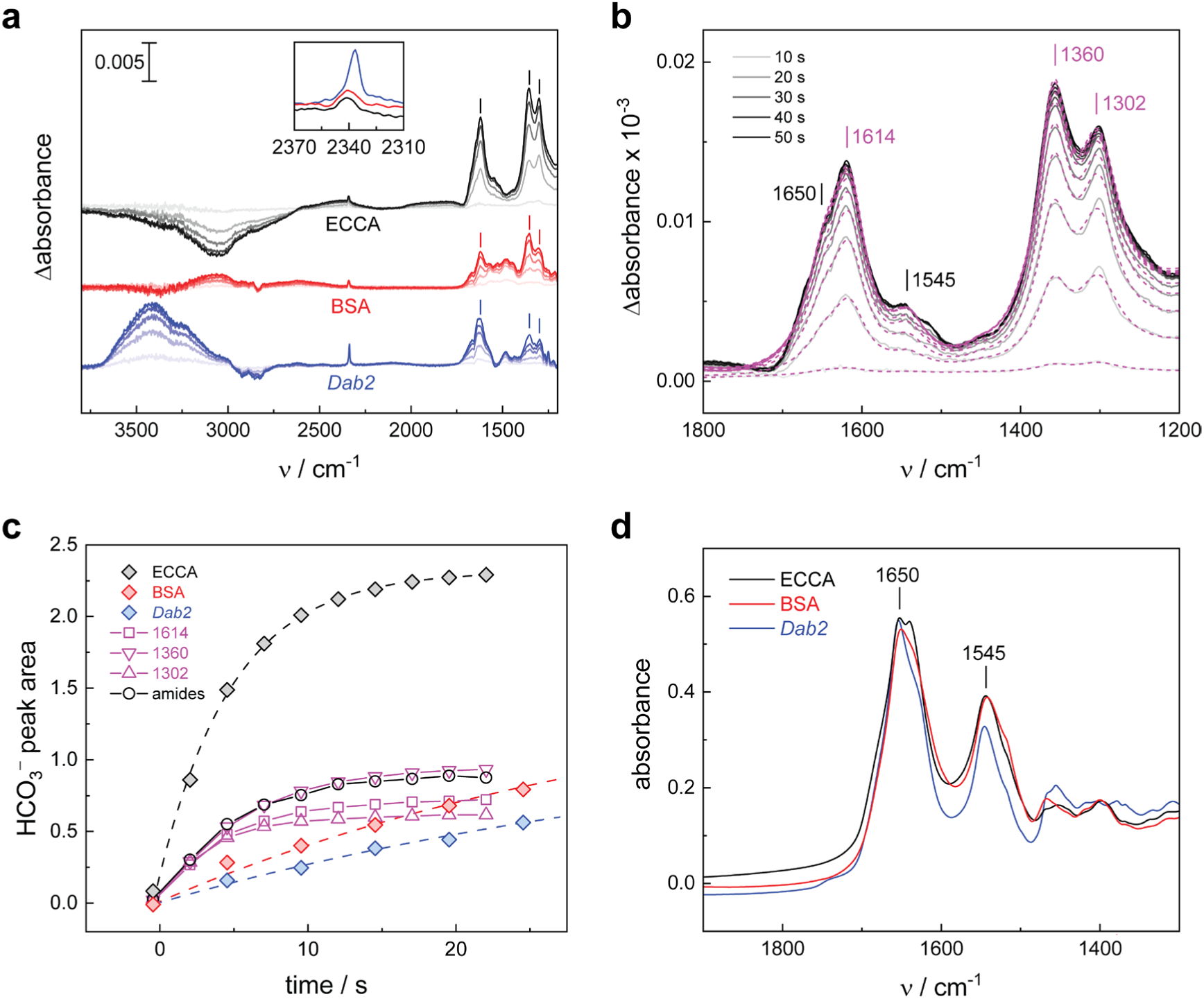
FTIR complete dataset. **a** Complete “CO_2_-minus-N_2_” difference spectra for ECCA, BSA, and *Dab2* between 3800 and 1200 cm^-1^. ECCA shows a characteristic consumption of H_2_O at approx. 3000 cm^-1^, as observed earlier^29^. The positive band at approx. 3400 cm^-1^ in the *Dab2* spectra hint at an unspecific increase of protein film hydration. The inset highlights the CO_2_ band as discussed in the main text (Fig. 3). **b** Exemplary fit of the ECCA difference spectra between 1800–1200 cm^-1^. The fit (dashed magenta traces) includes positive contributions from HCO_3_^−^ at 1614, 1360, and 1302 cm⁻¹ as well as the amide bands (compare panel **d**). **c** Time traces for the increase of HCO_3_^−^ in ECCA, BSA, and *Dab2* based on these fits. The peak area of the three components at 1614, 1360, and 1302 cm⁻¹ (shown explementary for ECCA, grey traces) are combined to one value (“HCO_3_^−^ peak area”) and plotted against time. While ECCA shows a fast logarithmic increase that reaches equilibrium after 20 s, BSA and Dab2 display a slow increase that fits the early, linear phase of product increase. **d** Comparison of the amide I (1650 cm^-1^) and amide II (1545 cm^-1^) bands of the ECCA, BSA, and *Dab2* protein films. The comparable amide ratio suggests similar hydration levels and protein concentrations in each film.

**Supplementary Figure 14.**
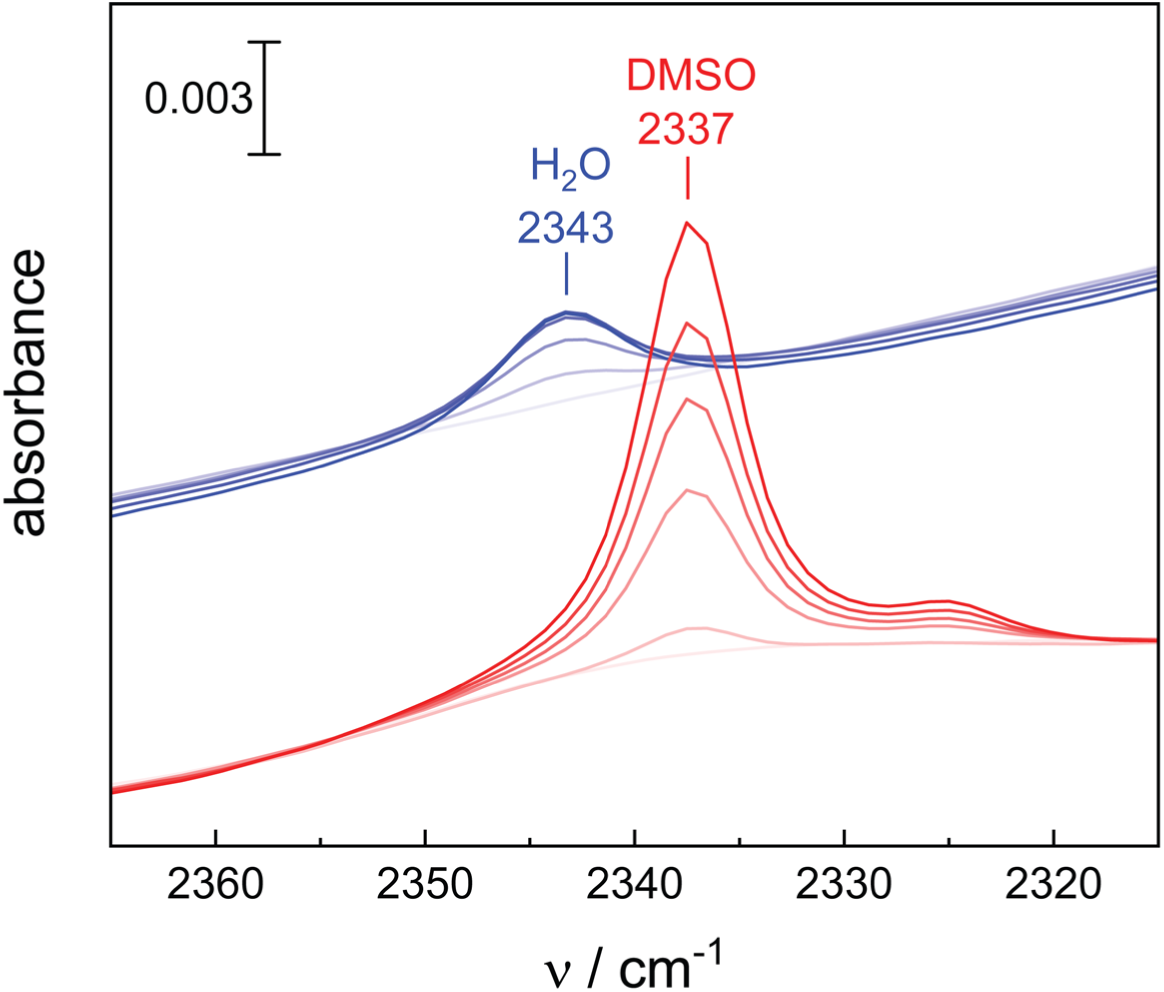
Absolute spectra of CO_2_ in solution with H_2_O or dimethyl sulfoxide (DMSO). Time traces for the increase of CO_2_ dissolved in H_2_O (blue traces) or DMSO (red traces) in the presence of 10% gaseous CO_2_. The 6 cm^-1^ difference in the CO_2_ marker band reflects the shift from a hydrogen-bonding, protic environment (H_2_O, 2342 cm^-1^) to a hydrophobic, aprotic environment (DMSO, 2337 cm^-1^).

**Supplementary Figure 15.**
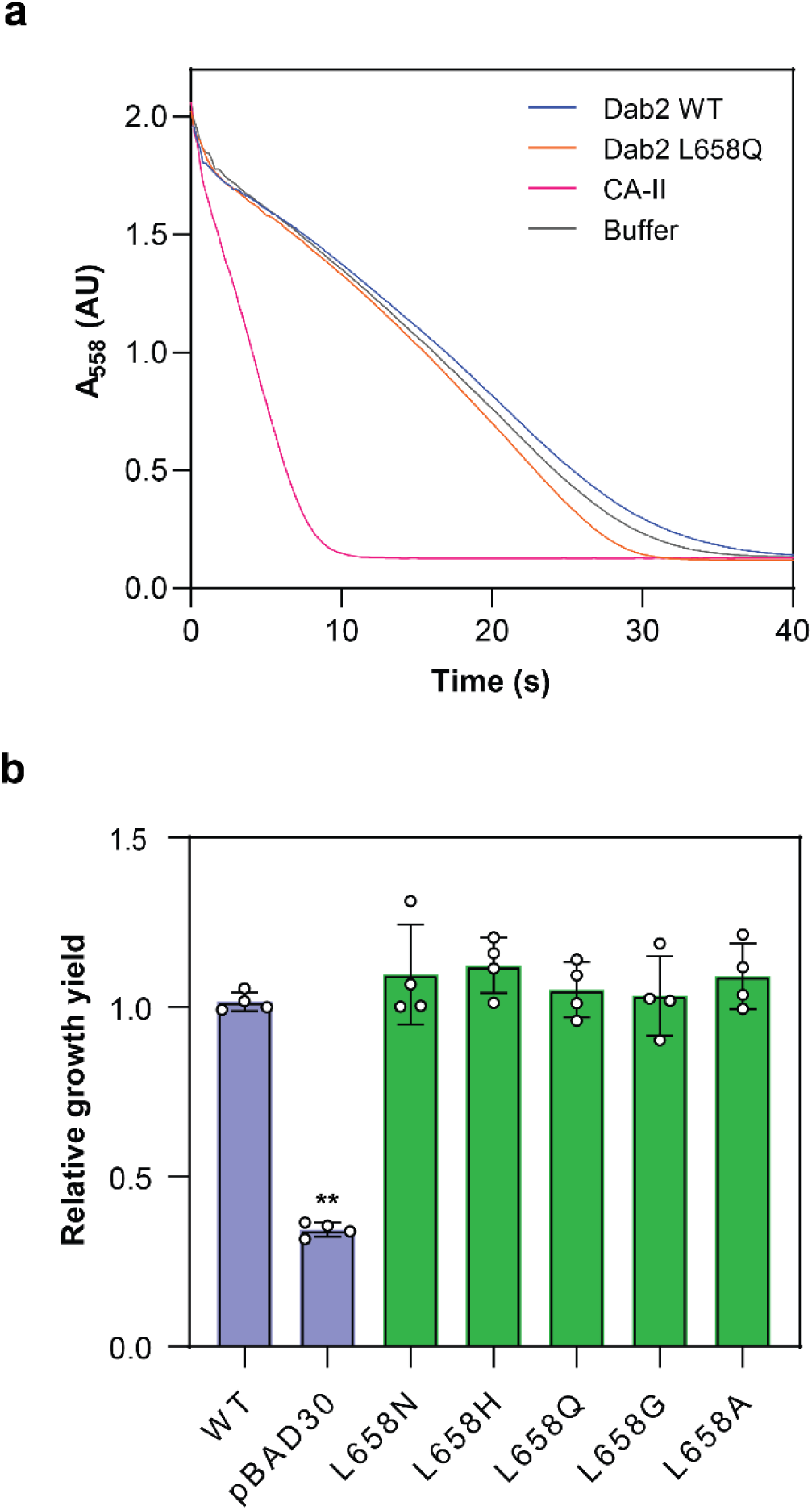
**Effect of L658 substitution on DAB2 activity**. **a** Representative results of CO_2_ hydration activity measured as a function of pH reduction, indicated by phenol red absorbance at 558 nm (n = 3 technical replicates). 5 nM Bovine carbonic anhydrase II (CA-II) was used as a positive control. Both Dab2 WT (500 nM) and L658Q variant (500 nM) showed similar background CO_2_ hydration as the negative control (Buffer). **b** Substitutions of Leu658 did not impair DAB2 ability to complement CA deficient *E. coli*. Bar heights and error bars represent means and standard deviations, respectively (n = 4 biological replicates). “**” Indicates statistically significant difference compared to WT (P < 0.05) according to Holm-Bonferroni corrected two-tailed t-test.

**Supplementary Figure 16.**
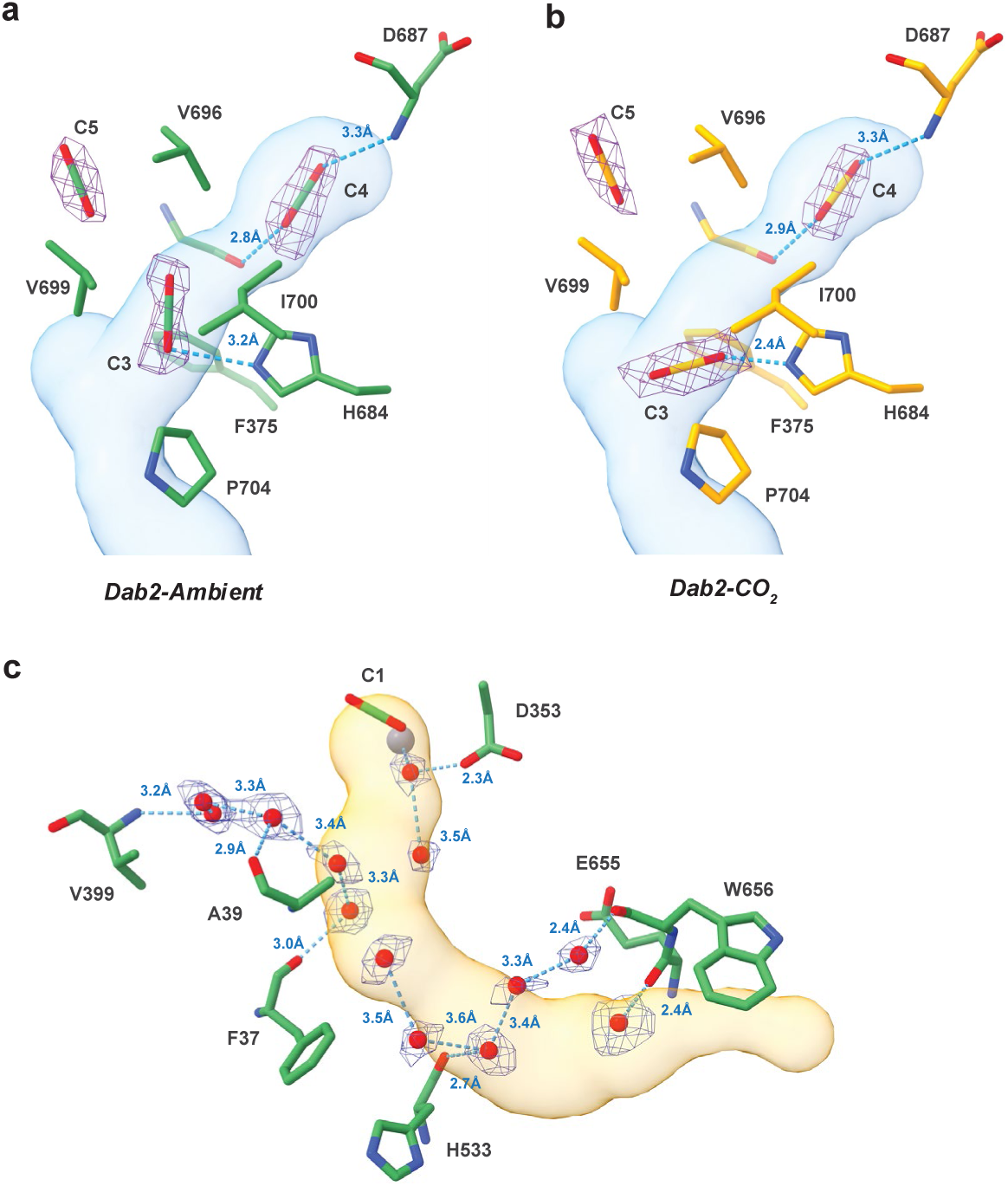
Ligands density fitting along the predicted tunnels. Coordination of CO_2_ molecules in **a** *Dab2-Ambient* and **b** *Dab2-CO_2_* within and near the T1 tunnel. Densities are shown at 6 σ. CO_2_ molecules are mainly stabilized by sidechain hydrophobic interactions. Blue dashes depict potential hydrogen-bonding with sidechain or backbone amide. **c** Coordination of *Dab2-Ambient* water molecules in the T2 tunnel. Densities of water molecules are displayed at 7 σ. Blue dash depicted the water molecule bonding network.

**Supplementary Figure 17.**
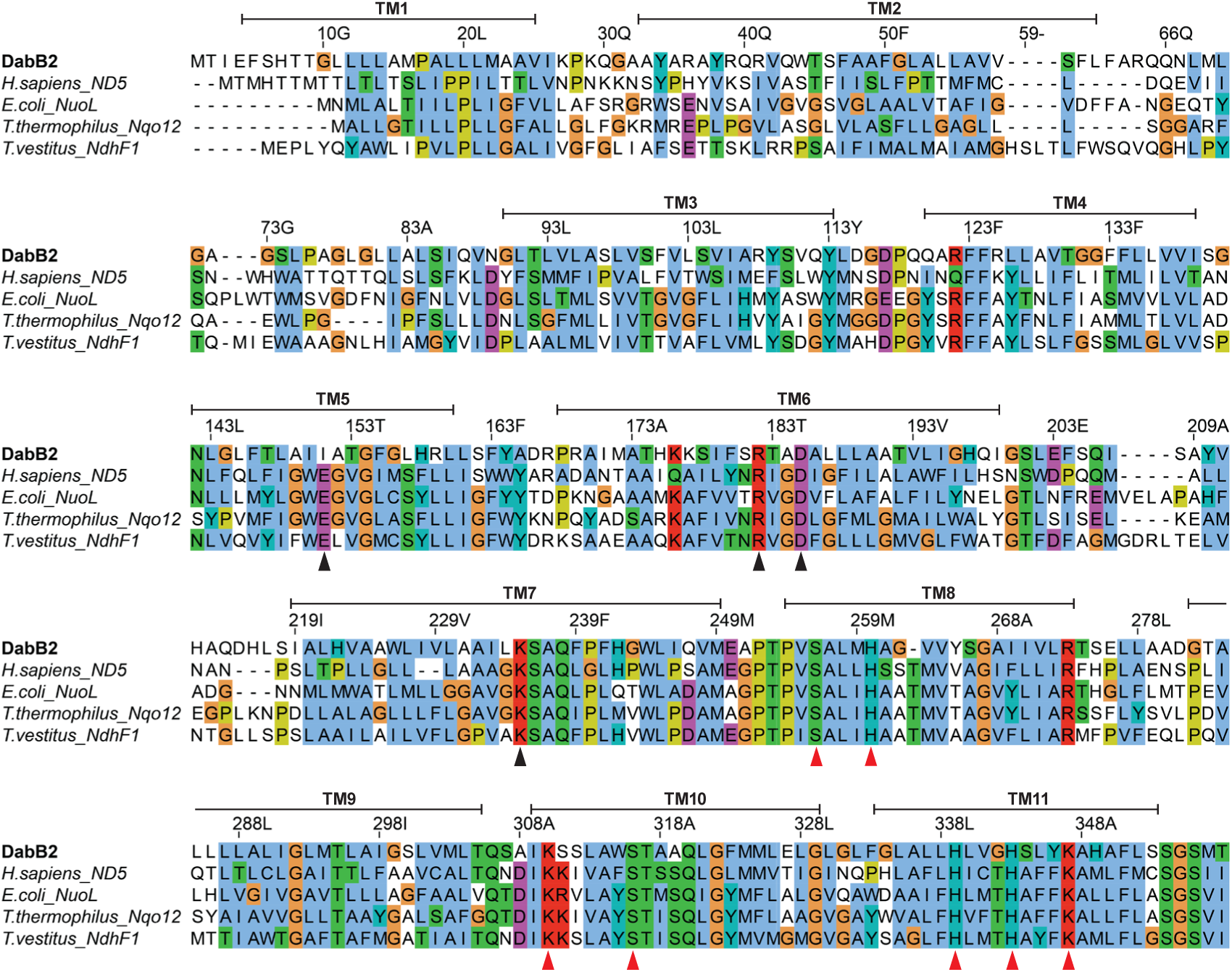
DabB2 Multiple sequence alignment. MUSCLE alignment of DabB2 transmembrane helixes TM1 to TM11 on Complex I (-like) distal proton-pumping subunits from *Homo sapiens*, *Escherichia coli*, *Thermus thermophilus* and *Thermosynechococcus vestitus*. Residues are colored by the *Clustal* coloring scheme. Regulatory ion-pairs and residues likely involved in proton transfer are marked by black and red triangle respectively. DabA2 residues number and secondary structure are shown above the alignment.

**Supplementary Figure 18.**
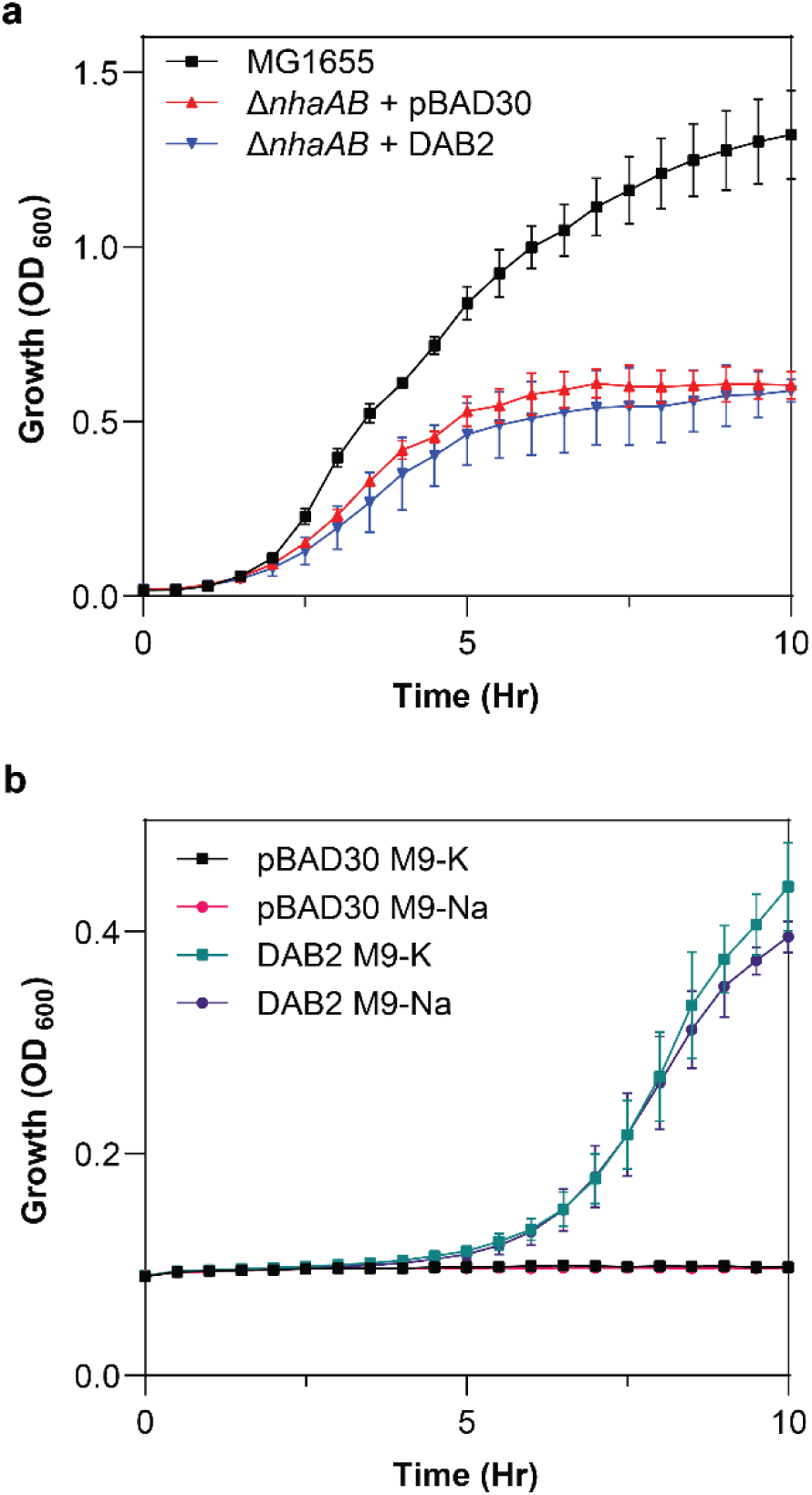
DAB2 activity is independent of sodium. **a** DAB2 did not restore growth of *E. coli* lacking both NhaA and NhaB sodium transporters under sodium stress (0.1 M). Wild-type *E. coli* MG1655 was used as a positive control. **b** Complementation of CA deficient *E. coli* by DAB2 in M9 medium prepared with potassium salts (M9-K) or sodium salts (M9-Na). The similar growth profile suggests the complex might not require sodium. Data points and error bars represent means and standard deviations, respectively (n = 4 biological replicates).

**Supplementary Figure 19.**
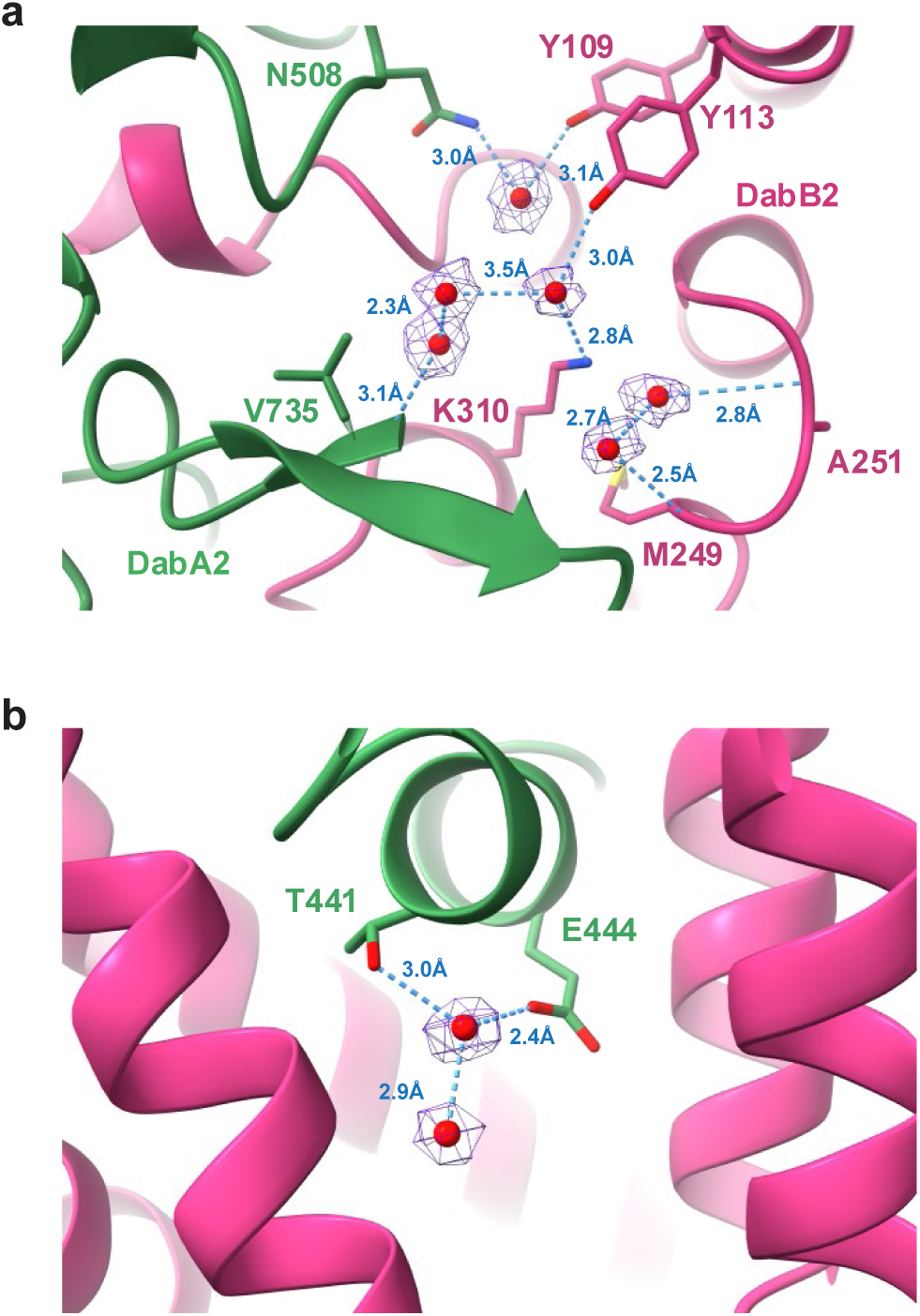
Putative proton pathway water molecules. Coordination of the water molecules shown in Fig. 5c at **a** the periplasmic side and **b** cytoplasmic side. Densities of water molecules are shown at 7 σ. Blue dashes depict hydrogen-bond with neighboring atoms.

## Notes

### Competing Interest Statement

The authors have declared no competing interest.

### Funding

- Deutsche Forschungsgemeinschaft

## References

1. Tcherkez, G. The mechanism of Rubisco-catalysed oxygenation. Plant, cell & environment 39, 983–997; 10.1111/pce.12629 (2016).

2. Maberly, S. C. & Gontero, B. Ecological imperatives for aquatic CO2-concentrating mechanisms. Journal of experimental botany 68, 3797–3814; 10.1093/jxb/erx201 (2017).

3. Raven, J. A. & Beardall, J. The ins and outs of CO 2. Journal of experimental botany 67, 1–13; 10.1093/jxb/erv451 (2016).

4. Price, G. D., Badger, M. R., Woodger, F. J. & Long, B. M. Advances in understanding the cyanobacterial CO2-concentrating-mechanism (CCM): functional components, Ci transporters, diversity, genetic regulation and prospects for engineering into plants. Journal of experimental botany 59, 1441–1461; 10.1093/jxb/erm112 (2008).

5. Singh, S. K., Sundaram, S. & Kishor, K. Carbon-Concentrating Mechanism. In Photosynthetic Microorganisms, edited by S. K. Singh, S. Sundaram & K. Kishor (Springer International Publishing, Cham, 2014), pp. 5–38.

6. Kupriyanova, E. V. et al. CO2-concentrating mechanism in cyanobacterial photosynthesis: organization, physiological role, and evolutionary origin. Photosynthesis research 117, 133–146; 10.1007/s11120-013-9860-z (2013).

7. Long, B. M., Rae, B. D., Rolland, V., Förster, B. & Price, G. D. Cyanobacterial CO2-concentrating mechanism components: function and prospects for plant metabolic engineering. Current opinion in plant biology 31, 1–8; 10.1016/j.pbi.2016.03.002 (2016).

8. Omata, T., Takahashi, Y., Yamaguchi, O. & Nishimura, T. Structure, function and regulation of the cyanobacterial high-affinity bicarbonate transporter, BCT1. Functional plant biology: FPB 29, 151–159; 10.1071/PP01215 (2002).

9. Price, G. D., Shelden, M. C. & Howitt, S. M. Membrane topology of the cyanobacterial bicarbonate transporter, SbtA, and identification of potential regulatory loops. Molecular membrane biology 28, 265–275; 10.3109/09687688.2011.593049 (2011).

10. Price, G. D., Woodger, F. J., Badger, M. R., Howitt, S. M. & Tucker, L. Identification of a SulP-type bicarbonate transporter in marine cyanobacteria. Proceedings of the National Academy of Sciences of the United States of America 101, 18228–18233; 10.1073/pnas.0405211101 (2004).

11. Shibata, M. et al. Distinct constitutive and low-CO2-induced CO2 uptake systems in cyanobacteria: genes involved and their phylogenetic relationship with homologous genes in other organisms. Proceedings of the National Academy of Sciences of the United States of America 98, 11789–11794; 10.1073/pnas.191258298 (2001).

12. Maeda, S.-I., Badger, M. R. & Price, G. D. Novel gene products associated with NdhD3/D4-containing NDH-1 complexes are involved in photosynthetic CO2 hydration in the cyanobacterium, Synechococcus sp. PCC7942. Molecular microbiology 43, 425–435; 10.1046/j.1365-2958.2002.02753.x (2002).

13. Mangiapia, M. et al. Proteomic and Mutant Analysis of the CO2 Concentrating Mechanism of Hydrothermal Vent Chemolithoautotroph Thiomicrospira crunogena. JOURNAL OF BACTERIOLOGY 199; 10.1128/JB.00871-16 (2017).

14. Scott, K. M. et al. Diversity in CO(2)-Concentrating Mechanisms among Chemolithoautotrophs from the Genera Hydrogenovibrio, Thiomicrorhabdus, and Thiomicrospira, Ubiquitous in Sulfidic Habitats Worldwide. Applied and environmental microbiology 85; 10.1128/AEM.02096-18 (2019).

15. Desmarais, J. J. et al. DABs are inorganic carbon pumps found throughout prokaryotic phyla. Nature microbiology 4, 2204–2215; 10.1038/s41564-019-0520-8 (2019).

16. Schmid, S. et al. Dissolved Inorganic Carbon-Accumulating Complexes from Autotrophic Bacteria from Extreme Environments. JOURNAL OF BACTERIOLOGY 203, e0037721; 10.1128/JB.00377-21 (2021).

17. Fan, S.-H. et al. MpsAB is important for Staphylococcus aureus virulence and growth at atmospheric CO2 levels. Nature communications 10, 3627; 10.1038/s41467-019-11547-5 (2019).

18. Fan, S.-H. et al. The MpsAB Bicarbonate Transporter Is Superior to Carbonic Anhydrase in Biofilm-Forming Bacteria with Limited CO 2 Diffusion. Microbiol Spectr 9; 10.1128/Spectrum.00305-21 (2021).

19. van Kempen, M. et al. Fast and accurate protein structure search with Foldseek. Nature biotechnology 42, 243–246; 10.1038/s41587-023-01773-0 (2024).

20. Holm, L., Laiho, A., Törönen, P. & Salgado, M. DALI shines a light on remote homologs: One hundred discoveries. Protein science: a publication of the Protein Society 32, e4519; 10.1002/pro.4519 (2023).

21. Rowlett, R. S. Structure and catalytic mechanism of the beta-carbonic anhydrases. Biochimica et biophysica acta 1804, 362–373; 10.1016/j.bbapap.2009.08.002 (2010).

22. Krissinel, E. & Henrick, K. Inference of Macromolecular Assemblies from Crystalline State. Journal of Molecular Biology 372, 774–797; 10.1016/j.jmb.2007.05.022 (2007).

23. Capasso, C. & Supuran, C. T. An overview of the alpha-, beta- and gamma-carbonic anhydrases from Bacteria: can bacterial carbonic anhydrases shed new light on evolution of bacteria? Journal of enzyme inhibition and medicinal chemistry 30, 325–332; 10.3109/14756366.2014.910202 (2015).

24. Smith, K. S., Ingram-Smith, C. & Ferry, J. G. Roles of the Conserved Aspartate and Arginine in the Catalytic Mechanism of an Archaeal β-Class Carbonic Anhydrase. J Bacteriol 184, 4240–4245; 10.1128/JB.184.15.4240-4245.2002 (2002).

25. Kimber, M. S. The active site architecture of Pisum sativum beta-carbonic anhydrase is a mirror image of that of alpha-carbonic anhydrases. The EMBO Journal 19, 1407–1418; 10.1093/emboj/19.7.1407 (2000).

26. Stripp, S. T. In Situ Infrared Spectroscopy for the Analysis of Gas-processing Metalloenzymes. ACS Catal. 11, 7845–7862; 10.1021/acscatal.1c00218 (2021).

27. Falk, M. & Miller, A. G. Infrared spectrum of carbon dioxide in aqueous solution. Vibrational Spectroscopy 4, 105–108; 10.1016/0924-2031(92)87018-B (1992).

28. Soboh, B. et al. The [NiFe]-hydrogenase accessory chaperones HypC and HybG of Escherichia coli are iron- and carbon dioxide-binding proteins. FEBS Letters 587, 2512–2516; 10.1016/j.febslet.2013.06.055 (2013).

29. Gomez, A. et al. Infrared spectroscopy reveals metal-independent carbonic anhydrase activity in crotonyl-CoA carboxylase/reductase. Chem. Sci. 15, 4960–4968; 10.1039/D3SC04208A (2024).

30. Sawaya, M. R. et al. The Structure of β-Carbonic Anhydrase from the Carboxysomal Shell Reveals a Distinct Subclass with One Active Site for the Price of Two. The Journal of biological chemistry 281, 7546–7555; 10.1074/jbc.M510464200 (2006).

31. Jin, S. et al. Structural insights into the LCIB protein family reveals a new group of β-carbonic anhydrases. Proc. Natl. Acad. Sci. U.S.A. 113, 14716–14721; 10.1073/pnas.1616294113 (2016).

32. Rowlett, R. S., Tu, C., Murray, P. S. & Chamberlin, J. E. Examination of the role of Gln-158 in the mechanism of CO(2) hydration catalyzed by beta-carbonic anhydrase from Arabidopsis thaliana. Archives of biochemistry and biophysics 425, 25–32; 10.1016/j.abb.2004.02.033 (2004).

33. Covarrubias, A. S. et al. Structure and Function of Carbonic Anhydrases from Mycobacterium tuberculosis. The Journal of biological chemistry 280, 18782–18789; 10.1074/jbc.M414348200 (2005).

34. Teng, Y.-B. et al. Structural insights into the substrate tunnel of Saccharomyces cerevisiae carbonic anhydrase Nce103. BMC structural biology 9, 67; 10.1186/1472-6807-9-67 (2009).

35. Alterio, V. et al. Zeta-carbonic anhydrases show CS2 hydrolase activity: A new metabolic carbon acquisition pathway in diatoms? Computational and structural biotechnology journal 19, 3427–3436; 10.1016/j.csbj.2021.05.057 (2021).

36. Chovancova, E. et al. CAVER 3.0: a tool for the analysis of transport pathways in dynamic protein structures. PLoS computational biology 8, e1002708; 10.1371/journal.pcbi.1002708 (2012).

37. Kaila, V. R. I., Wikström, M. & Hummer, G. Electrostatics, hydration, and proton transfer dynamics in the membrane domain of respiratory complex I. Proceedings of the National Academy of Sciences 111, 6988–6993; 10.1073/pnas.1319156111 (2014).

38. Kaila, V. R. I. Long-range proton-coupled electron transfer in biological energy conversion: towards mechanistic understanding of respiratory complex I. Journal of the Royal Society, Interface 15; 10.1098/rsif.2017.0916 (2018).

39. Di Luca, A., Gamiz-Hernandez, A. P. & Kaila, V. R. I. Symmetry-related proton transfer pathways in respiratory complex I. Proceedings of the National Academy of Sciences of the United States of America 114, E6314–E6321; 10.1073/pnas.1706278114 (2017).

40. Hoeser, F. et al. Respiratory complex I with charge symmetry in the membrane arm pumps protons. Proc. Natl. Acad. Sci. U.S.A. 119; 10.1073/pnas.2123090119 (2022).

41. Nakamaru-Ogiso, E. et al. The Membrane Subunit NuoL(ND5) Is Involved in the Indirect Proton Pumping Mechanism of Escherichia coli Complex I. The Journal of biological chemistry 285, 39070–39078; 10.1074/jbc.M110.157826 (2010).

42. Mühlbauer, M. E. et al. Water-Gated Proton Transfer Dynamics in Respiratory Complex I. Journal of the American Chemical Society 142, 13718–13728; 10.1021/jacs.0c02789 (2020).

43. Röpke, M., Saura, P., Riepl, D., Pöverlein, M. C. & Kaila, V. R. I. Functional Water Wires Catalyze Long-Range Proton Pumping in the Mammalian Respiratory Complex I. J. Am. Chem. Soc. 142, 21758–21766; 10.1021/jacs.0c09209 (2020).

44. Mayer, S., Steffen, W., Steuber, J. & Götz, F. The Staphylococcus aureus NuoL-Like Protein MpsA Contributes to the Generation of Membrane Potential. J Bacteriol 197, 794–806; 10.1128/JB.02127-14 (2015).

45. Denisov, I. G., Grinkova, Y. V., Lazarides, A. A. & Sligar, S. G. Directed Self-Assembly of Monodisperse Phospholipid Bilayer Nanodiscs with Controlled Size. J. Am. Chem. Soc. 126, 3477–3487; 10.1021/ja0393574 (2004).

46. Punjani, A., Rubinstein, J. L., Fleet, D. J. & Brubaker, M. A. cryoSPARC: algorithms for rapid unsupervised cryo-EM structure determination. Nature methods 14, 290–296; 10.1038/nmeth.4169 (2017).

47. Jumper, J. et al. Highly accurate protein structure prediction with AlphaFold. Nature 596, 583–589; 10.1038/s41586-021-03819-2 (2021).

48. Pettersen, E. F. et al. UCSF ChimeraX: Structure visualization for researchers, educators, and developers. Protein science: a publication of the Protein Society 30, 70–82; 10.1002/pro.3943 (2021).

49. Liebschner, D. et al. Macromolecular structure determination using X-rays, neutrons and electrons: recent developments in Phenix. *Acta crystallographica. Section D*, Structural biology 75, 861–877; 10.1107/S2059798319011471 (2019).

50. Emsley, P., Lohkamp, B., Scott, W. G. & Cowtan, K. Features and development of Coot. Acta crystallographica. Section D, Biological crystallography 66, 486–501; 10.1107/S0907444910007493 (2010).

51. Kunkel, T. A. Rapid and efficient site-specific mutagenesis without phenotypic selection. Proc. Natl. Acad. Sci. U.S.A. 82, 488–492; 10.1073/pnas.82.2.488 (1985).

52. Link, A. J., Phillips, D. & Church, G. M. Methods for generating precise deletions and insertions in the genome of wild-type Escherichia coli: application to open reading frame characterization. J Bacteriol 179, 6228–6237; 10.1128/jb.179.20.6228-6237.1997 (1997).

53. Harwood, C. R. & Cutting, S. M. Molecular biological methods for Bacillus / edited by Colin R. Harwood and Simon M. Cutting; with contributions by R. Chambert [and others] (Wiley, Chichester, 1990).

54. Wilbur, K. M. & Anderson, N. G. ELECTROMETRIC AND COLORIMETRIC DETERMINATION OF CARBONIC ANHYDRASE. The Journal of biological chemistry 176, 147–154; 10.1016/S0021-9258(18)51011-5 (1948).

55. Ashkenazy, H. et al. ConSurf 2016: an improved methodology to estimate and visualize evolutionary conservation in macromolecules. Nucleic acids research 44, W344–50; 10.1093/nar/gkw408 (2016).

56. Waterhouse, A. M., Procter, J. B., Martin, D. M. A., Clamp, M. & Barton, G. J. Jalview Version 2--a multiple sequence alignment editor and analysis workbench. Bioinformatics (Oxford, England) 25, 1189–1191; 10.1093/bioinformatics/btp033 (2009).

57. Schada von Borzyskowski, L., et al. Implementation of the β-hydroxyaspartate cycle increases growth performance of Pseudomonas putida on the PET monomer ethylene glycol. Metabolic engineering 76, 97–109; 10.1016/j.ymben.2023.01.011 (2023).

58. Demichev, V., Messner, C. B., Vernardis, S. I., Lilley, K. S. & Ralser, M. DIA-NN: neural networks and interference correction enable deep proteome coverage in high throughput. Nature methods 17, 41–44; 10.1038/s41592-019-0638-x (2020).

59. Glatter, T. et al. Large-scale quantitative assessment of different in-solution protein digestion protocols reveals superior cleavage efficiency of tandem Lys-C/trypsin proteolysis over trypsin digestion. Journal of proteome research 11, 5145–5156; 10.1021/pr300273g (2012).

60. Ahrné, E., Molzahn, L., Glatter, T. & Schmidt, A. Critical assessment of proteome-wide label-free absolute abundance estimation strategies. Proteomics 13, 2567–2578; 10.1002/pmic.201300135 (2013).

61. Oliveira, S. H. P. et al. KVFinder: steered identification of protein cavities as a PyMOL plugin. BMC bioinformatics 15, 197; 10.1186/1471-2105-15-197 (2014).

